# Post-translational modifications orchestrate the intrinsic signaling bias of orphan receptor GPR52

**DOI:** 10.1101/2024.10.26.620377

**Authors:** Bingjie Zhang, Mengna Ma, Wei Ge, Shanshan Li, Guang Yang, Huilan Wang, Boxun Lu, Wenqing Shui

## Abstract

Despite recent advances in GPCR structure and pharmacology, the regulation of GPCR activation, signaling, and functioning by diverse post-translational modifications (PTMs) has been largely unexplored. Furthermore, for a special category of self-activating orphan GPCRs, it is completely unknown whether and how specific PTMs control their unique signaling profiles and cellular functions. In this study of GPR52, an orphan GPCR with exceedingly high constitutive G protein signaling activity and emerging as a neurotherapeutic target, we discovered its disproportionately low arrestin recruitment activity. Through profiling the N-glycosylation and phosphorylation patterns of the receptor and clarifying their roles in G protein *versus* arrestin coupling, we found these two types of PTMs in specific motifs differentially shape the intrinsic signaling bias of GPR52. While N-terminal N-glycosylation promotes constitutive Gs signaling possibly through favoring the self-activating conformation, phosphorylation in helix 8, to our great surprise, suppresses arrestin recruitment and thus attenuates receptor internalization. In addition, we uncovered the counteracting roles of N-glycosylation and phosphorylation in modulating GPR52-dependent accumulation of the huntingtin protein (HTT) in brain striatal cells. Thus, our study provides novel insights into the regulation of intrinsic signaling bias and cellular function of an orphan GPCR *via* distinct PTMs in different motifs.

## Introduction

G protein-coupled receptors (GPCRs), the largest family of cell surface receptor proteins in mammals, transduce a vast diversity of signals into appropriate cellular responses and control a broad range of physiological processes. Despite the diversity of extracellular ligands and physiological roles of GPCRs, these cell surface receptors share a conserved molecular architecture and intracellular transducers^1-3^. Agonist binding stabilizes active conformations of the receptor, facilitating the binding of one or more cytosolic transducers including the heterotrimeric G proteins that dissociate to α and βγ subunits upon activation to initiate downstream signaling^1,3^. Activated GPCRs can also recruit arrestin proteins whose primary role is to block G protein coupling and facilitate receptor internalization^4^. However, arrestins also function as scaffold proteins for the initiation of additional signaling that seems to be G protein-independent^4,5^. Compared to a pathway-balanced reference ligand, a number of agonists are known to elicit stronger signaling through one pathway over another, for example favoring G protein over arrestin coupling or vice versa, which is termed ligand-dependent signaling bias^1,6,7^. Therapeutic exploitation of signaling bias is expected to increase drug efficacy while avoiding adverse effects attributable to particular pathways^1,8,9^.

Unlike agonist-induced activation, there exists a special category of GPCRs that are constitutively activated without any external ligand stimulation. Most of these self-activating GPCRs such as GPR52, GPR21, GPR3, and the adhesion GPCR family are orphan receptors with unknown endogenous ligands^10^. Notably, the constitutive activity of these receptors refers to their high basal activity of Gs protein coupling to elicit downstream cAMP signaling, for which the structural mechanism has been revealed using ligand-free receptor bound to heterotrimeric Gs protein^10-15^. However, for these self-activating receptors with high constitutive G protein activity, it remains elusive whether their arrestin pathway possesses a balanced or unbalanced activity.

Post-translational modifications (PTMs) are known to regulate GPCR signaling dynamics and physiological responses as well as control receptor biogenesis and trafficking^16,17^. For example, glycosylation naturally occurring in the biosynthetic pathway in general controls GPCR folding, maturation, and transport to the cell surface^18-20^. Agonist activation of a GPCR typically induces phosphorylation in the cytoplasmic C-terminus and loop regions, which promotes arrestin recruitment and activation^2^. Despite the general knowledge, the effects of different PTMs on individual receptor activation and downstream pathway coupling as well as underlying molecular mechanisms are poorly understood for the majority of GPCRs^16,18^. Furthermore, for these self-activating GPCRs, it is completely unknown whether and how specific PTMs regulate their unique signaling profiles and cellular functions.

Here, we used GPR52 as an exemplar GPCR to characterize the influences of PTMs on its pathway-specific coupling activity and cellular function. GPR52 is a brain-enriched orphan receptor with exceedingly high constitutive Gs-coupled activity^10,11^. It has emerged as a promising therapeutic target for the treatment of Huntington’s disease (HD), schizophrenia, and other psychiatric disorders^21^. Particularly, GPR52 is reported to regulate the accumulation of mutant huntingtin protein (mHTT) in striatal neurons of HD animal models^22-24^. Previous structural elucidation revealed its unique self-activating mechanism, and yet no PTM moieties were observed in the receptor structures due to their high conformational flexibility^11^.

In this study, we first discovered the disproportionately low constitutive arrestin recruitment activity of GPR52 compared to its high basal Gs signaling activity. The weak β-arrestin activity was not much elevated by stimulation with a surrogate agonist. This self-driven preference of the Gs pathway over the arrestin pathway by GPR52 is termed intrinsic signaling bias, a paradigm that has been rarely explored. We further investigated the underlying mechanisms from the perspective of site-specific or motif-specific PTMs, and characterized relevant functional impact on huntingtin protein regulation in a cellular HD model.

## Results

### Intrinsic signaling bias and blocked internalization of GPR52

We conducted several cellular activity assays in HEK293T cells transfected with a given receptor to assess the strengths of activating different pathways by GPR52 in comparison to GLP-1R. GLP-1R is a prototypical class B GPCR known to exhibit very low basal activity, and its endogenous agonist GLP-1(7-36) induces balanced signaling between Gs and β-arrestin pathways^25^. As expected, GLP-1R without agonist stimulation showed no measurable activity in both Gs-mediated cAMP accumulation and Gαs-γ1 dissociation measured by bioluminescent resonance energy transfer^26^, and reached 100% full response upon induction by a saturating amount of GLP-1(7-36) (Figures 1A-1D). In contrast, GPR52 at the unliganded state displayed unusually high basal Gs activity. Relative to the full response induced by a potent synthetic agonist wo-459 (EC_50_ = 30.6 nM), unliganded GPR52 achieved 83.4% and 87.4% maximal efficacy (*E*_max_) in the cAMP accumulation and Gs dissociation assays, respectively (Figures 1A-1D). This result confirmed the robust constitutive Gs-coupled activity of GPR52 which is marginally increased by agonist stimulation.

**Figure 1.**
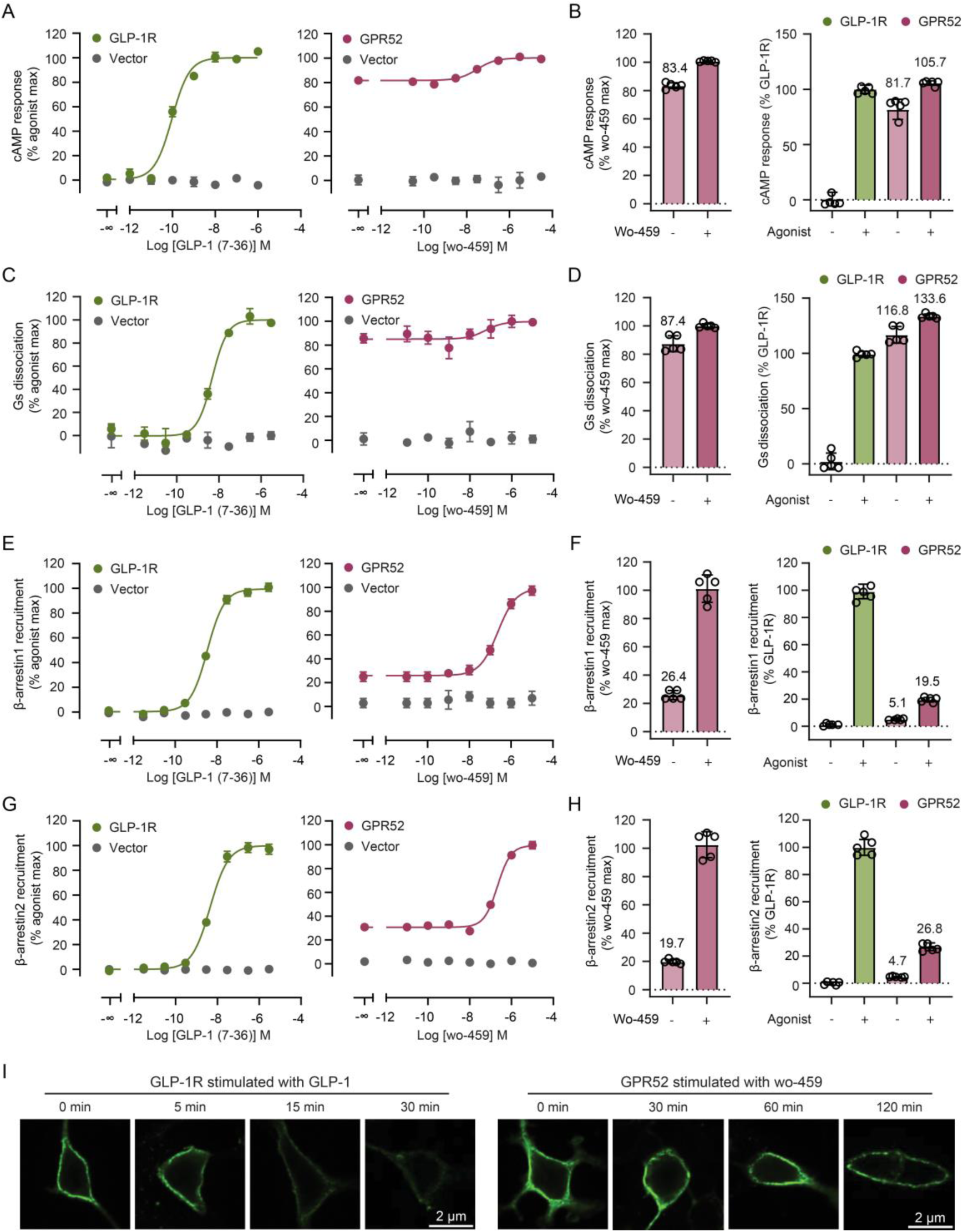
Intrinsic signaling bias and blocked internalization of GPR52 expressed in HEK293T cells. (A) Constitutive activity and agonist-induced activity of GLP-1R (left) and GPR52 (right) characterized by the cAMP accumulation assay. (B) Maximal efficacy in the cAMP accumulation assay normalized to the agonist-induced full response of GPR52 (left) or to the agonist-induced full response of GLP-1R (right). (C, D) Activity profiles same as (A, B) characterized by the Gs dissociation assay. (E. F) Activity profiles same as (A, B) characterized by the BRET-based β-arrestin1 recruitment assay. (G, H) Activity profiles same as (A, B) characterized by the BRET-based β-arrestin2 recruitment assay. In (A-H), data are represented as mean ± s.d. (bars) from five independent experiments. Agonist EC_50_ values determined from three independent experiments are summarized in Table S1. (I) Immunofluorescence imaging of HEK293T cells expressing FLAG-tagged GLP-1R and treated with 1 μM GLP-1 for different time points (left), and cells expressing FLAG-tagged GPR52 and treated with 10 μΜ wo-459 for different time points (right).

Surprisingly, when we measured the β-arrestin recruitment activity of GPR52 still normalized by the agonist-induced full response of the respective pathway, we found unliganded GPR52 only displayed 26.4% *E*_max_ and 19.7% *E*_max_ in recruiting β-arrestin1 and β-arrestin2, respectively (Figures 1E-1H). As the above activity measurement was based on BRET assays, we then conducted an orthogonal Tango assay to show unliganded GPR52 exhibited 39.2% *E*_max_ relative to the agonist-induced full response (Supplementary Figures 1A, 1B). Therefore, compared to the substantially high constitutive Gs signaling activity (83.4-87.4% *E*_max_ in two independent assays), GPR52 displays a disproportionately low constitutive β-arrestin recruitment activity (19.7-39.2% *E*_max_ in two independent assays).

Noticing that the above constitutive activity was normalized to the full response of GPR52 itself, we then compared the pathway activation strengths between GPR52 and GLP-1R. Setting the agonist-induced full response of GLP-1R as the reference, we observed 81.7% and 116.8% *E*_max_ achieved by unliganded GPR52 and 105.7% and 133.6% *E*_max_ by agonist-induced GPR52 in activating cAMP accumulation and Gs dissociation, respectively (Figures 1B, 1D). It indicates that GPR52 is able to self-activate the Gs pathway to an extent near or above the maximal efficacy of agonist-induced GLP-1R. In contrast, in the BRET-based β-arrestin recruitment assays, while unliganded GPR52 showed 5.1% and 4.7% *E*_max_ in recruiting β-arrestin1 and β-arrestin2 relative to the GLP-1R full response, agonist induction only moderately increased the β-arrestin activity (19.5% *E*_max_ for β-arrestin1, 26.8% *E*_max_ for β-arrestin2). In these assays, *E*_max_ measurement was not affected by receptor surface expression which was comparable for GPR52 and GLP-1R (Supplementary Figures 1C-1E). Thus, GPR52 does not self-activate the β-arrestin pathway as robustly as the Gs pathway. In fact, its β-arrestin recruitment activity with or without agonist stimulation was significantly lower than that of a prototypical GPCR induced by a full agonist.

Given that β-arrestin recruitment is critical for initiating receptor internalization, we sought to examine the internalization behavior of GPR52 *vs* GLP-1R. While 5-min agonist treatment induced evident internalization of GLP-1R from the plasma membrane and more intense internalization was observed at 15 min treatment, GPR52 invariably remained on the cell surface without appreciable internalization in the absence or presence of agonist for as long as 120 min (Figure 1I, Supplementary Figures 2A-2D). The almost undetectable internalization of GPR52 with or without agonist stimulation corroborates its relatively weak β-arrestin recruitment activity.

In summary, using multiple pharmacological assays, our study recapitulates the high constitutive Gs signaling activity of GPR52 and discovers its low constitutive β-arrestin recruitment activity. Furthermore, compared to agonist-induced GLP-1R signaling efficacy, agonist-treated GPR52 also exhibits robust Gs coupling and weak β-arrestin coupling, leading to the blockage of receptor internalization. To substantiate this finding, we measured GPR52 activities with another synthetic agonist EX5467, and observed the same magnitude of β-arrestin1/2 activity attenuation and Gs activity enhancement compared to agonist-induced GLP-1R (Supplementary Figures 3A-3H).

Therefore, unlike conventional ligand bias, this preferential activation of the Gs pathway over the β-arrestin pathway by GPR52 is more likely to be a ligand-independent system bias arising from the receptor’s inherent properties. Thus, it is referred to as the intrinsic signaling bias of GPR52.

### N-terminal N-glycosylation promotes constitutive Gs signaling and cell surface expression of GPR52

Intrigued by the question of whether and how post-translational modifications contribute to the intrinsic signaling bias of GPR52, we set out to investigate the role of N-linked glycosylation which predominantly occurs at the N-terminal ectodomain and extracellular loops (ECLs) of many mammalian GPCRs^16-18^. Notably, folded ECL2 in the unliganded GPR52 structure is reported to act as a built-in agonist which confers the receptor with a high basal activity in Gs signaling^11^. Although we did not find any N-glycosylation consensus sites (N-X-S/T) in ECL2, we did find three of them in the N-terminus (N2, N13, N20) of the GPR52 protein sequence (Figure 2A). The receptor exogenously expressed in HEK293T cells showed multiple bands designating its monomer and aggregated forms in the cell lysate. Most of these bands were down-shifted following the peptide-N-glycosidase (PNGase F) treatment, indicating the receptor underwent N-glycosylation (Figure 2B). Site-directed mutagenesis of three potential N-glycosylation sites (Asn to Gln) resulted in moderate down-shifting of major protein bands for all three single-site mutants (N2Q, N13Q, and N20Q) compared to the wild-type, and the triple mutant 3NxQ displayed the most significant down-shifting movement (Figure 2C). These results suggested that GPR52 expressed in HEK293T cells carries N-glycosylation possibly at three consensus sites in the N-terminus.

**Figure 2.**
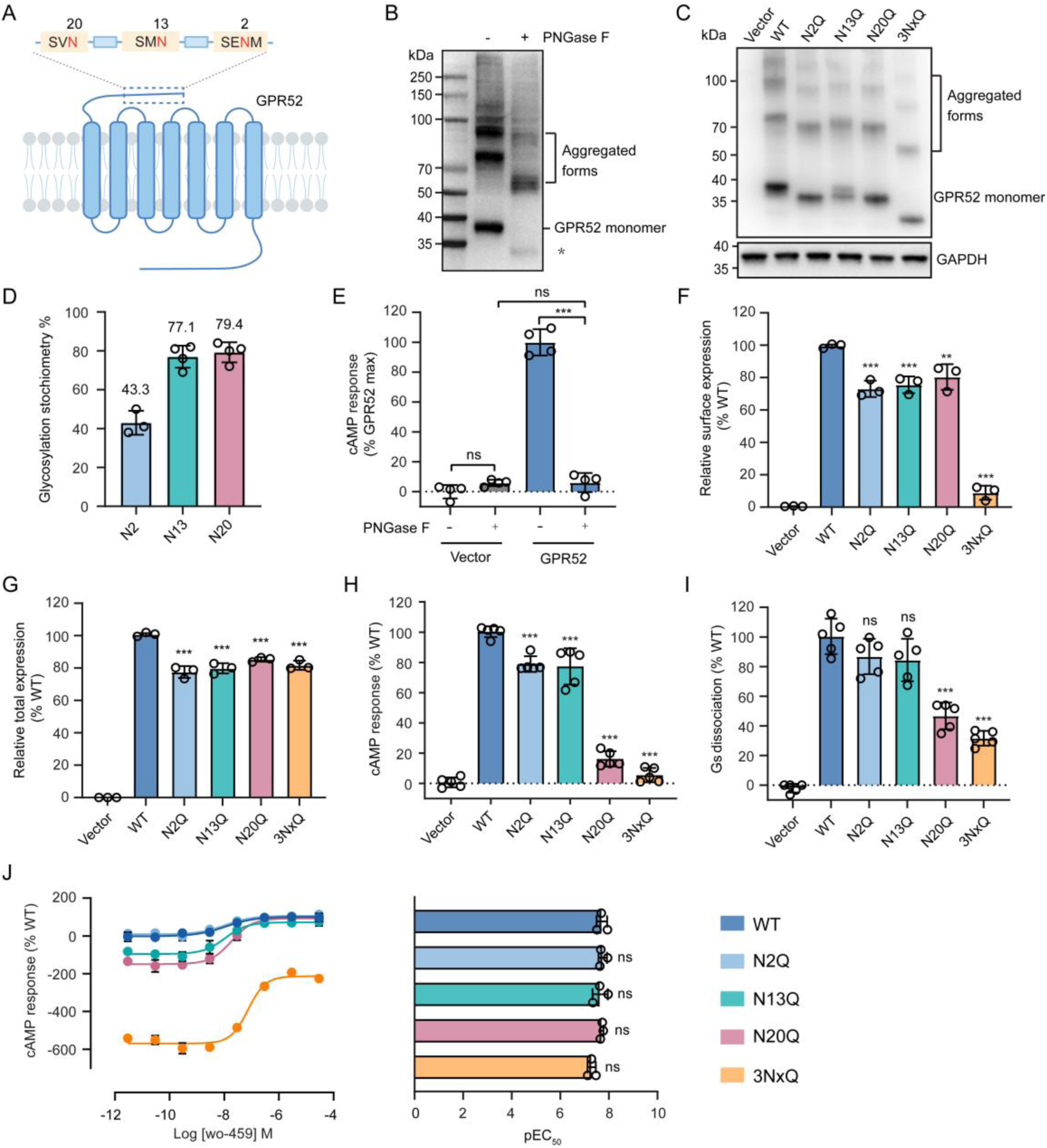
N-terminal N-glycosylation contributes to GPR52-mediated Gs signaling and receptor surface expression. (A) Putative N-glycosylation sites (shown in red) in the consensus motif N-X-S/T (X≠P) at the N-terminus of GPR52. (B) Western blot of FLAG-tagged GPR52 protein expressed in HEK293T cells treated with (+) or without (-) PNGase F. The asterisk indicates the deglycosylated form of GPR52. (C) Western blot of FLAG-tagged wild-type (WT), single mutants (N2Q, N13Q, N20Q), and triple mutant (3NxQ) of GPR52 expressed in HEK293T cells. (D) N-glycosylation stoichiometry at specific Asn residues of GPR52 measured by quantitative MS analysis. Data are represented as mean ± s.d. from at least three independent experiments. (E) Constitutive cAMP accumulation measured in GPR52-expressing HEK293T cells treated with or without PNGase F. Data are represented as mean ± s.d. from at least four independent experiments. (F, G) Cell surface (F) and total (G) expression levels of HA-tagged WT and N-glycosylation mutants of GPR52 expressed in HEK293T cells (for the cAMP accumulation assay) determined by flow cytometry. Data are represented as mean ± s.d. from three independent experiments. (H, I) The effects of N-glycosylation site-directed mutagenesis of GPR52 on the constitutive Gs activity measured by cAMP accumulation (H) and Gs dissociation (I) assays in HEK293T cells. Data are represented as mean ± s.d. from five independent experiments. (G) Concentration-response curves (left) and potency (right) of agonist wo-459 in activating WT or N-glycosylation mutants of GPR52 measured by the cAMP accumulation assay. Data are shown as mean ± s.d. from three independent experiments. Statistics significance was assessed by one-way ANOVA with Dunnett’s multiple comparison test. \*\**p* < 0.01, \*\*\**p* < 0.001, and ns, no significance.

To directly verify the N-glycosylation sites, we purified recombinant GPR52 protein with an intact N-terminus fused with a BRIL for stabilization^11^ (Supplementary Figure 4A). Mass spectrometry analysis of the protein digest unambiguously pinpointed N-glycosylation occurring on three consensus sites (Supplementary Figures 4B-4D), and quantification of the glycosylated and unglycosylated peptides containing the same sites allowed us to estimate glycosylation stoichiometry at specific sites (43.3% at N2, 77.1% at N13, and 79.4% at N20) (Figure 2D). Therefore, we identified site-specific N-glycosylation at the N-terminus with varying abundance.

To investigate the functional impact of N-glycosylation on GPR52-mediated Gs signaling, we treated receptor-expressing HEK293T cells with PNGase F and measured cAMP accumulation with or without agonist simulation. Interestingly, the constitutive Gs signaling activity was almost completely abolished for unliganded GPR52 after deglycosylation (Figure 2E), and yet the agonist wo-459 was able to activate both the glycosylated and deglycosylated receptor with the same potency (Supplementary Figures 5A, 5B). Given that PNGase F treatment also caused ∼40% reduction of GPR52 surface expression (Supplementary Figure 5C), we conclude that enzymatic depletion of N-glycosylation impairs both receptor trafficking to the cell surface and constitutive Gs signaling.

With regard to N-glycosylation single and triple mutants, the substitution of all three modification sites resulted in substantially reduced surface expression of GPR52 (8.9% and 6.0% in the cAMP accumulation and Gs dissociation assays respectively relative to 100% of wild-type) whereas single mutants at different sites retained the majority of surface expression (73%-80% in the cAMP accumulation assay, 57%-64% in the Gs dissociation assay) (Figure 2F, Supplementary Figure 5D). All tested mutants displayed total protein expression comparable to wild-type (Figure 2G, Supplementary Figure 5D). Remarkably, the Gs signaling activity of unliganded mutant N20Q dropped to 16.3% *E*_max_ in the cAMP accumulation assay and 46.7% *E*_max_ in the Gs dissociation assay relative to the full response of wild-type GPR52. In comparison, Gs activities of the other two mutants N2Q and N13Q were marginally decreased or unaffected in two assays (Figures 2H, 2I). Considering that the surface expression of mutant N20Q was only reduced by 20% and GPR52 was found to maintain cAMP accumulation to the maximal efficacy even with a loss of 60% surface expression (Supplementary Figure 5E), we speculated that the dramatic drop of constitutive Gs activity for mutant N20Q was primarily attributed to its deficient N-glycosylation rather than reduced surface expression. As for the triple mutant 3NxQ, however, significant attenuation of its constitutive Gs activity mainly resulted from the almost complete loss of surface expression (Figure 2F).

Agonist stimulation of three single mutants including N20Q further activated the receptor to elevate Gs-mediated cAMP accumulation to the maximal efficacy of wild-type with similar potency. In addition, the agonist was able to activate the triple mutant in Gs signaling with the same potency as wild-type (Figure 2J).

Taken together, our study indicates that N-glycosylation at residue N20 strongly promotes constitutive Gs signaling of GPR52 yet has little effect on agonist-induced Gs activity. Moreover, N-glycosylation at three N-terminal residues collectively contributes to receptor trafficking to the cell surface.

### Site-specific N-glycosylation favors the formation of GPR52 self-activating conformation

A previous study reveals unique structural characteristics of GPR52, which involves ECL2 occupying the orthosteric binding pocket and serving as a built-in agonist to confer the high basal Gs activity^11^ (Figure 3A). Our discovery of the role of N-glycosylation in mediating constitutive Gs signaling of GPR52 prompted us to investigate whether site-specific N-glycosylation influences the conformational features of the unliganded receptor.

**Figure 3.**
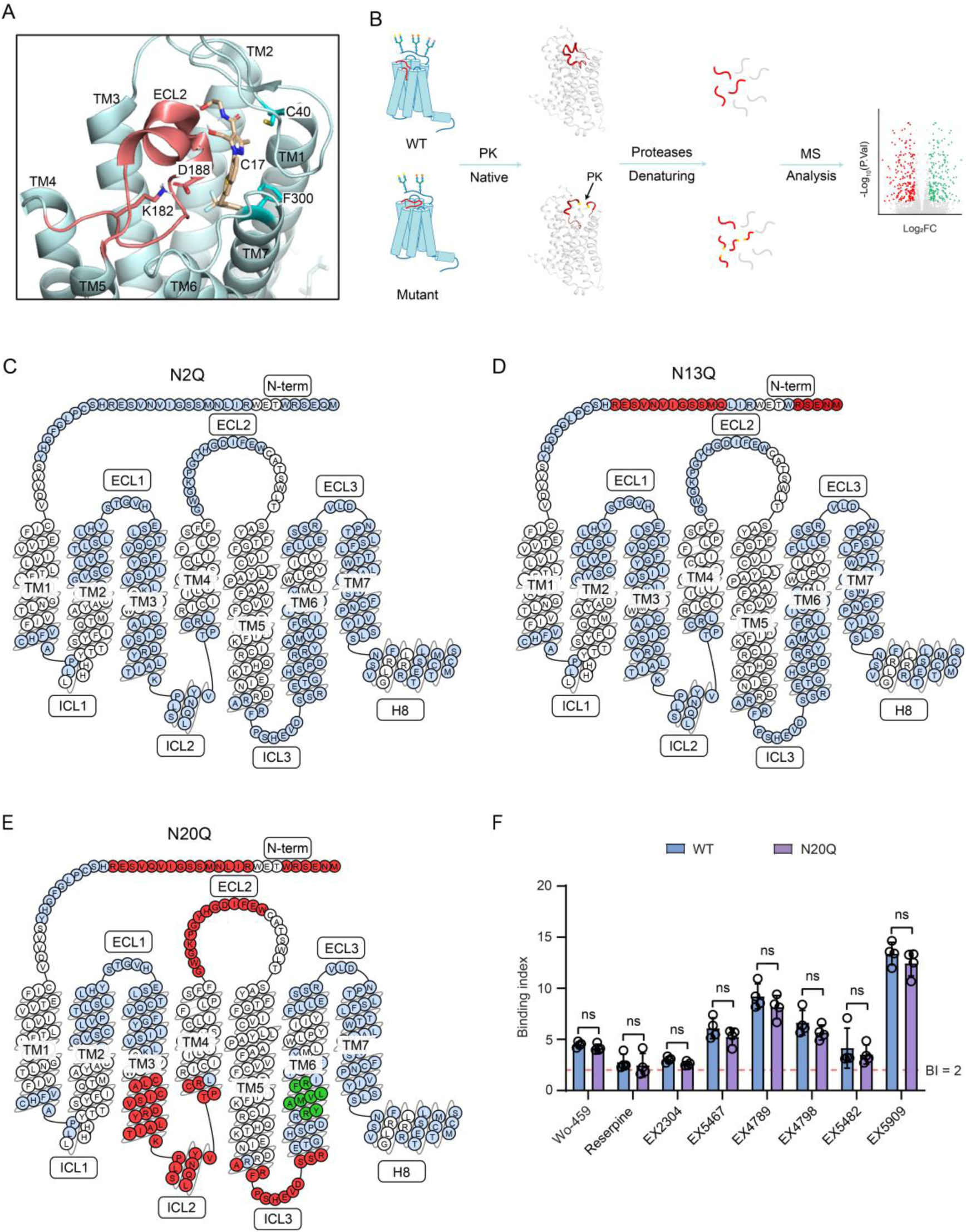
Effects of site-specific N-glycosylation on GPR52 native conformation and agonist binding property. (A) Structural view of ECL2 (shown in red) occupying the orthosteric pocket and agonist c17 (shown in wheat) located in the side pocket of GPR52 (PDB code 6LI0). (B) Schematic of the LiP-MS workflow for assessing protein conformational alterations. Purified apo GPR52 protein of wild-type and mutants were treated with proteinase K (PK) in a native buffer, followed by proteolysis with proteases (trypsin and chymotrypsin) under a denaturing condition, to generate tryptic peptides for MS analysis. Differential tryptic peptides between a mutant and the wild-type indicate conformational changes of specific regions^28^. (C-E) LiP-MS derived proteolytic pattern of a given GPR52 mutant relative to wild-type displayed in the snake plot from GPCRdb^51^. Red or green residues from tryptic peptides with differential abundances between the mutant and wild-type indicate structurally altered regions in the mutant (red, increased PK accessibility; green, decreased PK accessibility). Blue residues from unchanged tryptic peptides between the mutant and wild-type indicate unaltered structural regions. Quantitative LiP-MS data for each mutant *vs* WT is summarized in Table S2. (F) Binding index (BI) of each GPR52 agonist to purified GPR52 WT or N20Q protein measured by the affinity MS assay. An average BI > 2 (*p* < 0.05) indicate positive ligand binding to the protein^31,32^. Data are shown as mean ± s.d. from four independent experiments. Statistical significance was assessed by two-tailed Student’s t-test. ns, no significance.

Here we adapted the limited proteolysis-mass spectrometry (LiP-MS) approach to probing the conformational changes of GPR52 as a result of N-glycosylation site-directed mutagenesis. LiP-MS has become a powerful technology to monitor proteome-wide protein structural changes by measuring the variation of protein susceptibility to a promiscuous protease^27-30^. In brief, wild-type and three N-glycosylation site mutants of GPR52 with an intact N-terminus were purified in vitro (Supplementary Figures 6A-6E). Each protein was subjected to limited proteolysis by a nonspecific protease (proteinase K, PK) in the native buffer, followed by complete proteolysis with dual proteases (trypsin and chymotrypsin) in a denaturing condition to yield a tryptic digest amenable to bottom-up MS analysis^28^ (Figure 3B, see details in Methods). Tryptic peptides with differential abundances between each mutant and the wild-type arise from varying susceptibility of the respective structural region to PK cleavage, thus indicating local conformational alterations between conditions^28^.

In comparison to the wild-type, the N2Q and N13Q mutants showed proteolytic patterns with no or very few structurally altered regions (Figures 3C, 3D). In contrast, the N20Q mutant generated a much more disrupted proteolytic pattern, with significant reduction of tryptic peptides yielded from the N-terminus, ECL2, the intracellular half of TM3, ICL2, and ICL3 (Figure 3E). Importantly, this result indicates that depletion of N-glycosylation at N20 may cause increased conformational flexibility of ECL2, which could lead to disruption of its agonist-like motif that is required for receptor self-activation. In addition, as both the cytoplasmic end of TM3 and ICL2 are engaged in the Gs binding cavity^11^, their conformational changes in the N20Q mutant relative to wild-type indicated by LiP-MS analysis substantiates the impaired Gs signaling activity of the mutant. In summary, the LiP-MS result supports the notion that N-glycosylation at N20, but not two other glycosylation sites, could facilitate the formation of GPR52 native conformation (especially in ECL2 and Gs binding site) that promotes receptor self-activation and Gs coupling.

Given that a synthetic agonist capable of further increasing the Gs activity of GPR52 is located in an allosteric side pocket close to the extracellular surface^11^ (Figure 3A), we then assessed the effect of site-specific N-glycosylation on ligand binding to the receptor. Using the affinity mass spectrometry approach developed in our lab to detect ligand binding to purified GPCR proteins^31-34^, we compared the binding profiles of a series of agonists to the N20Q mutant *vs* wild-type GPR52. All agonists showed comparable binding indexes between the two conditions, suggesting that deglycosylation at N20 does not affect ligand binding to the side pocket (Figure 3F). Altogether, these insights into receptor conformational and ligand binding attributes corroborate the finding from our cellular signaling data that N-glycosylation at N20 potentiates the constitutive Gs activity yet not the agonist-induced Gs activity of GPR52.

### Profiling intracellular phosphorylation patterns of ligand-free and agonist-induced GPR52

Upon agonist-induced activation, GPCRs are routinely bound and phosphorylated at Ser and Thr residues at the cytoplasmic side by specific GPCR kinases (GRKs) or effector kinases. Receptor phosphorylation is a key functional PTM to determine the subsequent binding of arrestin proteins^35-38^. Because of the strong self-activation of GPR52, we suspect the receptor may undergo phosphorylation at an unliganded state. To test this hypothesis, we first purified the apo GPR52 protein and performed MS analysis of the protein digest for direct profiling of phosphorylation sites (Figure 4A). To increase our chance of mapping all possible phosphorylation events, we also expressed full-length GPR52 in HEK293T cells and enriched phosphopeptides from the protein digestion products prior to MS analysis (Figure 4B). From the unenriched sample, we identified five phosphorylation sites in ICL3 or helix 8 while phospho-enrichment allowed us to capture two additional phosphosites in helix 8 and the C-terminal tail (C-tail) (Figure 4C). Notably, site-specific phosphorylation stoichiometry measurement based on MS quantification revealed a rather high phosphorylation level at S332 (18.8%) in helix 8 together with two other sites of lower phosphorylation levels (T334, 3.9%; S338, 3.4%) in the same motif (Figure 4D). MS spectra for peptides bearing phosphorylation at S332 or S338 are shown in Figure 4E, and spectra for other identified phosphopeptides are summarized in Supplementary Figure 7. As virtually all GPCR phosphorylation is reported to occur within the C-tail or on the ICLs^16,17^, the presence of multiple phosphosites in helix 8 for unliganded GPR52 may be associated with its unique self-activation and signaling mechanism.

**Figure 4.**
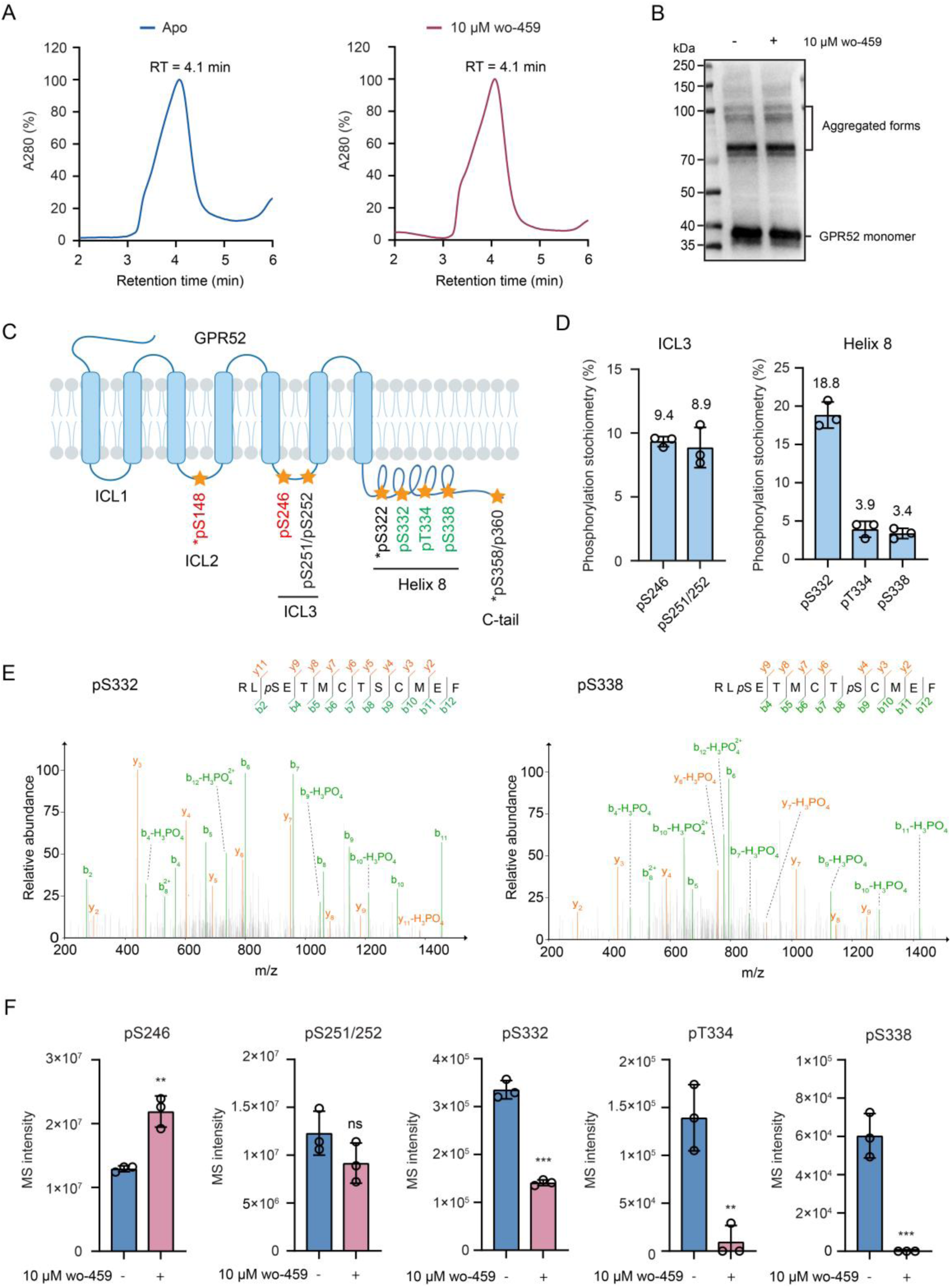
Intracellular phosphorylation profiles of GPR52 in the absence or presence of agonist stimulation. (A) Analytical size-exclusion chromatograms of purified GPR52 protein expressed in HEK293F cells with or without 10 μM wo-459 treatment. (B) Western blot of full-length HA-tagged GPR52 expressed in HEK293T cells with or without 10 μM wo-459 treatment. (C) Schematic of identified phosphorylation sites in different intracellular motifs of GPR52 with or without wo-459 treatment. Under the agonist treatment, phosphosites that appeared or showed increased phosphorylation are highlighted in red, those disappeared or showed reduced phosphorylation are in green, and those with no phosphorylation changes are in black. Asterisks indicate phosphosites only detected in phospho-enrichment samples. (D) Phosphorylation stoichiometry at specific residues of apo GPR52 protein measured by quantitative MS analysis. (E) Representative MS/MS spectra for GPR52 peptides bearing phosphorylation at S332 or S338. (F) Relative changes of site-specific phosphorylation levels in ICL3 or helix 8 of GPR52 isolated from agonist-treated cells compared to untreated cells. Phosphorylation levels are reflected by MS intensity of respective phosphopeptides detected at specific conditions. Data are shown as mean ± s.d. from three independent experiments. Statistics significance was assessed by two-tail Student’s t-test. \*\**p* < 0.01, \*\*\**p* < 0.001, and ns, no significance.

Next, we profiled the phosphorylation pattern of GPR52 upon stimulation with a saturating amount of agonist using both the purified recombinant protein and the full-length protein expressed in the cells (Figures 4A, 4B). Surprisingly, quantitative comparison indicates that phosphorylation at three residues in helix 8 (S332, T334, S338) was either attenuated or completely abolished upon synthetic agonist treatment whereas phosphorylation at two residues in ICL3 (S246, S251/S252) was either increased or unchanged (Figure 4F). Additionally, a new phosphosite S148 was identified in ICL2 through phospho-enrichment analysis of the agonist-induced GPR52 (Figure 4C). The unexpected finding of robust basal phosphorylation in helix 8 (H8) and its attenuation in agonist-induced GPR52 prompted us to look into the functional impact of phosphorylation on the arrestin pathway.

### Helix 8 phosphorylation suppresses arrestin coupling and attenuates internalization of GPR52

To clarify the role of motif-specific phosphorylation in mediating the arrestin pathway activity, we made Ala substitutions on all identified phosphosites within each motif, thus generating ICL2-A, ICL3-A, H8-A, and C tail-A mutants (Figure 5A). An additional site-specific mutant (S332A in H8) was prepared considering its highest basal phosphorylation level. As β-arrestin2 is more likely to be associated with GPR52 signaling *in vivo*^39^, we first assessed the constitutive β-arrestin2 recruitment activity of these mutants. Relative to unliganded wild-type, the ICL2-A mutant showed reduced activity with 43.9% *E*_max_ in the Tango assay whereas the ICL3-A mutant displayed no change of activity in both Tango and BRET assays. Conversely, the H8-A mutant substantially elevated its constitute β-arrestin2 recruitment activity to 300% *E*_max_ and 451% *E*_max_ relative to wild-type in Tango and BRET assays, respectively (Figure 5B). Similarly, both the C tail-A and S332A mutants showed increased β-arrestin2 recruitment activity, though to a lesser extent than the activity of the H8-A mutant (Figure 5B). To support the regulatory role of motif-specific phosphorylation, we then made phosphomimic substitution (Ser/Thr to Asp) on respective phosphosites to generate ICL2-D and H8-D mutants (Figure 5A). Consistently, the H8-D mutant showed significantly impaired constitute β-arrestin2 recruitment activity compared to the H8-A mutant in both assays and to the wild type in the Tango assay (Figure 5C). The ICL2-D mutant did not show an opposite effect to the ICL2-A mutant, which is conceivable due to the complicated influence of mutagenesis (Figure 5C).

**Figure 5.**
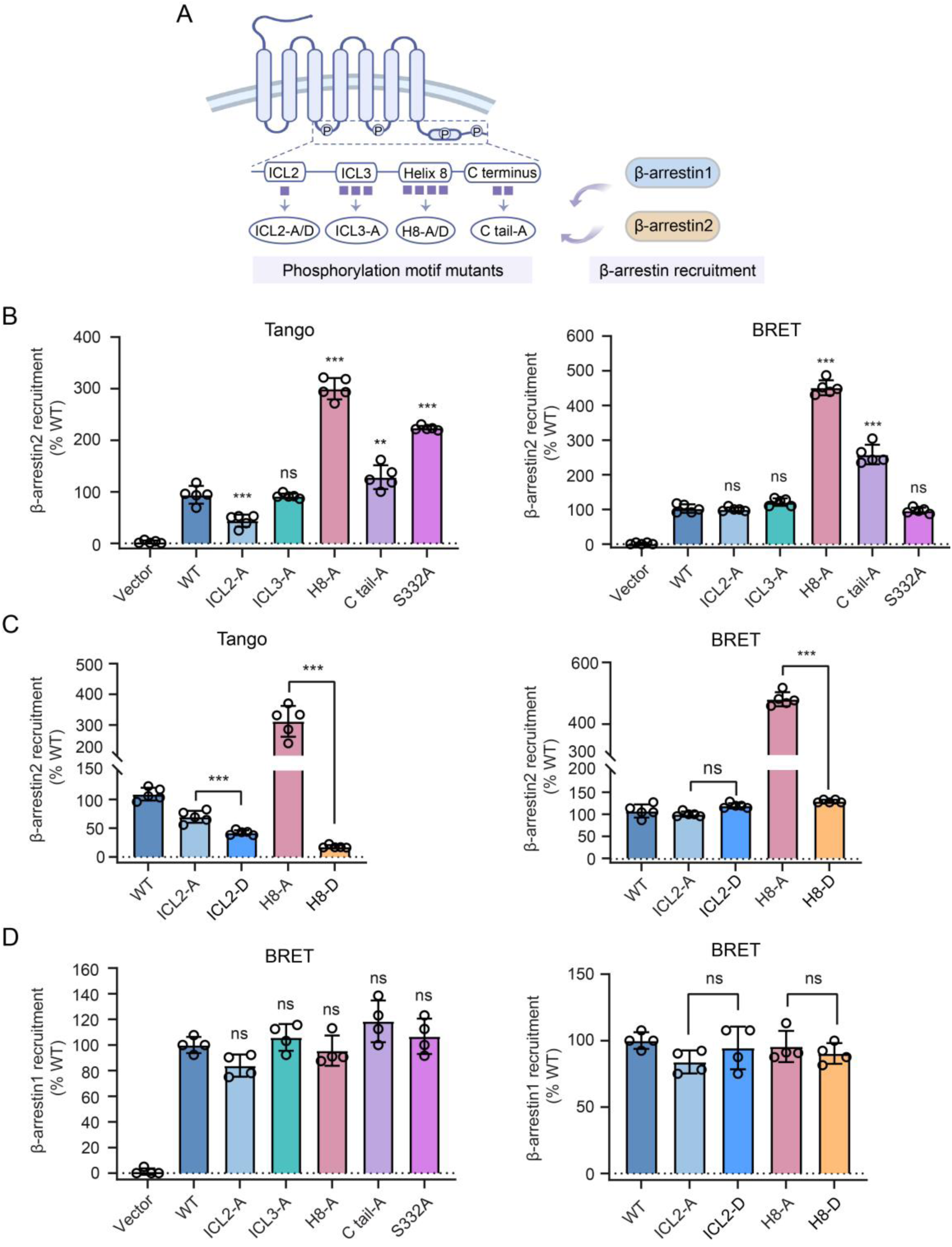
Function of motif-specific phosphorylation in mediating GPR52 constitutive activity of arrestin recruitment. (A) Design of phosphorylation mutants in different GPR52 motifs for assessing arrestin recruitment activity in HEK293T cells. (B) Constitutive β-arrestin2 recruitment activity of GPR52 phospho-deficient mutants relative to WT measured by Tango (left) and BRET (right) assays. Data are represented as mean ± s.d. from five independent experiments. (C) Constitutive β-arrestin2 recruitment activity of GPR52 phosphomimic mutants relative to WT measured by Tango (left) and BRET (right) assays. Data are represented as mean ± s.d. from five independent experiments. (D) Constitutive β-arrestin1 recruitment activity of GPR52 phospho-deficient (left) and phosphomimic (right) mutants relative to WT measured by BRET assays. Data are shown as mean ± s.d. from four independent experiments. Statistics significance was assessed by one-way ANOVA with Dunnett’s multiple comparison test. \*\**p* < 0.01, \*\*\**p* < 0.001 and ns, no significance.

In summary, these results suggest that phosphorylation at S148 in ICL2 likely facilitates the constitutive β-arrestin2 recruitment to GPR52, and yet unexpectedly, phosphorylation in helix 8 plays an autoinhibitory role in constitutive β-arrestin2 recruitment. The inhibitory effect of helix 8 is in part contributed by S332 phosphorylation. Of note, none of these mutations significantly affected cell surface expression of GPR52, indicating varied β-arrestin2 recruitment was not strong enough to perturb basal receptor internalization which was kept low for mutants as much as for wild-type (Supplementary Figures 8A, 8B). Furthermore, none of these motif- or site-specific mutants showed significant changes in their constitutive activity in recruiting β-arrestin1 (Figure 5D), highlighting the selective role of phosphorylation in mediating engagement of the more physiologically relevant β-arrestin2 transducer.

Next, to examine the contribution of helix 8 phosphorylation to agonist-induced arrestin recruitment, we measured the dose-response activity of H8 mutants and wild-type. Remarkably, compared to the agonist-induced full response of wild-type, the H8-A mutant achieved an efficacy as high as 332% *E*_max_ and 421% *E*_max_ in agonist-induced β-arrestin2 recruitment determined by Tango and BRET assays, respectively (Figures 6A, 6B). In contrast, the H8-D mutant showed an opposite effect by suppressing the agonist-induced β-arrestin2 recruitment activity in both assays (Figures 6A, 6B). Interestingly, although the H8-A mutant had little effect on constitutive β-arrestin1 recruitment, its activity in recruiting β-arrestin1 upon agonist stimulation was considerably increased to 470% *E*_max_ relative to wild-type in the BRET assay (Figure 6C). Moreover, mutation of the phosphosites in helix 8 primarily affected the maximal efficacy and yet hardly changed the potency of agonist-induced β-arrestin coupling (Figure 6D).

**Figure 6.**
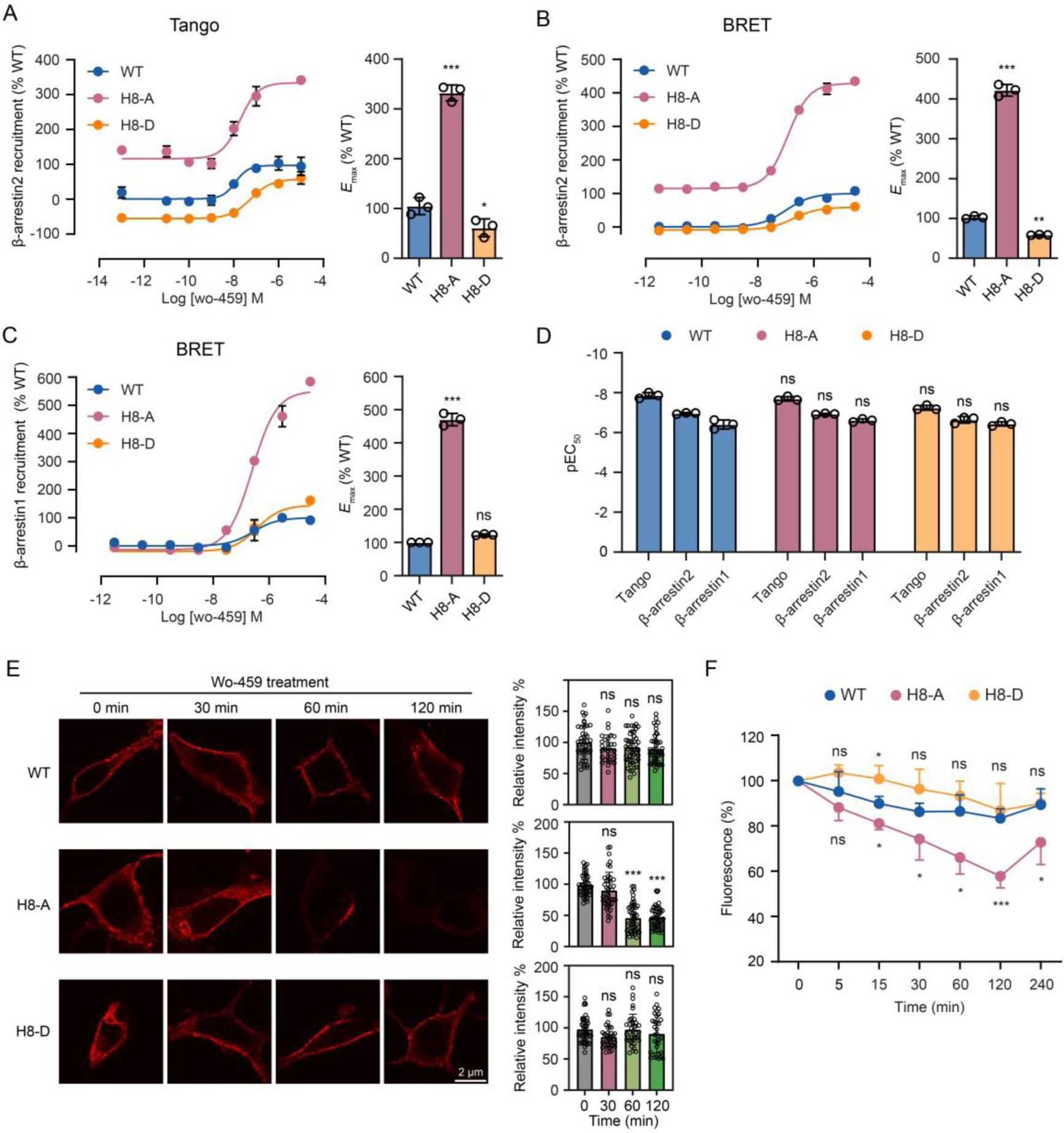
Effects of helix 8 phosphorylation on agonist-induced arrestin recruitment activity and internalization of GPR52. (A, B) Concentration-response curves (left) and efficacy (right) of agonist wo-459-induced β-arrestin2 recruitment activity of WT, H8-A and H8-D mutants of GPR52 expressed in HEK293T cells determined by Tango (A) and BRET (B) assays. (C) Concentration-response curves (left) and efficacy (right) of agonist wo-459-induced β-arrestin1 recruitment activity of WT, H8-A and H8-D mutants of GPR52 expressed in HEK293T cells determined by the BRET assay. In (A-C), data are represented as mean ± s.d. from three independent experiments. (D) Potency of agonist wo-459 in activating β-arrestin recruitment to WT, H8-A and H8-D mutants of GPR52 related to data in (A-C). Agonist EC_50_ values are summarized in Table S1. Data are represented as mean ± s.d. from three independent experiments. (E) Representative immunofluorescence imaging (left) and fluorescence intensity quantification (right) of WT, H8-A and H8-D mutants of HA-tagged GPR52 expressed in HEK293T cells treated with 10 μM wo-459 for different incubation times. Quantification data are represented as mean ± s.e.m (n ≧ 31 cells). (F) Internalization of WT, H8-A and H8-D mutants of HA-tagged GPR52 expressed in HEK293T cells treated with 10 μM wo-459 for different incubation times measured by flow cytometry. Data are represented as mean ± s.d. from four independent experiments. Statistics significance was assessed by one-way ANOVA with Dunnett’s multiple comparison test. \**p* < 0.05, \*\**p* < 0.01, \*\*\**p* < 0.001 and ns, no significance.

As we uncovered the tremendous enhancement of β-arrestin2 and β-arrestin1 recruitment to the H8-A mutant, we wondered whether such an effect would alter agonist-induced receptor internalization which is rather weak for wild-type GPR52 as a result of its intrinsic signaling bias. By immunofluorescence, we observed a significant loss of receptor surface staining at 60 min post-agonist treatment for the H8-A mutant, yet not for the wild-type or the H8-D mutant (Figure 6E). More quantitative analysis by flow cytometry reveals enhanced agonist-induced internalization of the H8-A mutant at 15 min to 240 min post-ligand treatment whereas wild-type and the H8-D mutant exhibited almost no internalization over the same period (Figure 6F, Supplementary Figure 9A). Therefore, we conclude that helix 8 phosphorylation attenuates both constitutive and agonist-induced receptor internalization possibly through inhibiting arrestin coupling.

### N-glycosylation and phosphorylation shape the intrinsic signaling bias of GPR52 and modulate HTT levels in a cellular HD model

To explore how PTMs influence the constitutive coupling profile of GPR52, we measured the basal activity of both N-glycosylation and phosphorylation mutants on the alternative pathways. As expected, three N-glycosylation single-site mutants showed reduced constitutive β-arrestin2 recruitment activity relative to wild-type, with the most profound reduction observed on the N20Q mutant, which is associated with their impaired Gs signaling activity (Supplementary Figure 10A). On the other side, mutation of the phosphosite on ICL2 had little influence on constitutive Gs activity of GPR52 whereas that on helix 8 moderately impaired Gs activity (Supplementary Figure 10B). Combining all cellular activity data, we plotted the constitutive coupling strengths of Gs and arrestin pathways for PTM mutants and wild-type based on their % *E*_max_ measurement relative to the agonist-induced full response of wild-type (Figure 7A). All PTM mutants shown here have surface expression comparable to the wild-type in respective assays (Supplementary Figures 10C, 10D). Similar to the wild-type, the N-glycosylation mutants N2Q and N13Q, and phosphorylation mutants ICL2-A/D and H8-D, all display an intrinsic bias towards Gs signaling. In contrast, the H8-A mutant shows an opposite bias towards arrestin coupling, and the N20Q mutant is coupled to both severely impaired pathways (Figure 7A). Thus, N-glycosylation at N20 and phosphorylation in helix 8 both shape the intrinsic signaling bias of GPR52, which would presumably affect the receptor-mediated function in the native tissue.

**Figure 7.**
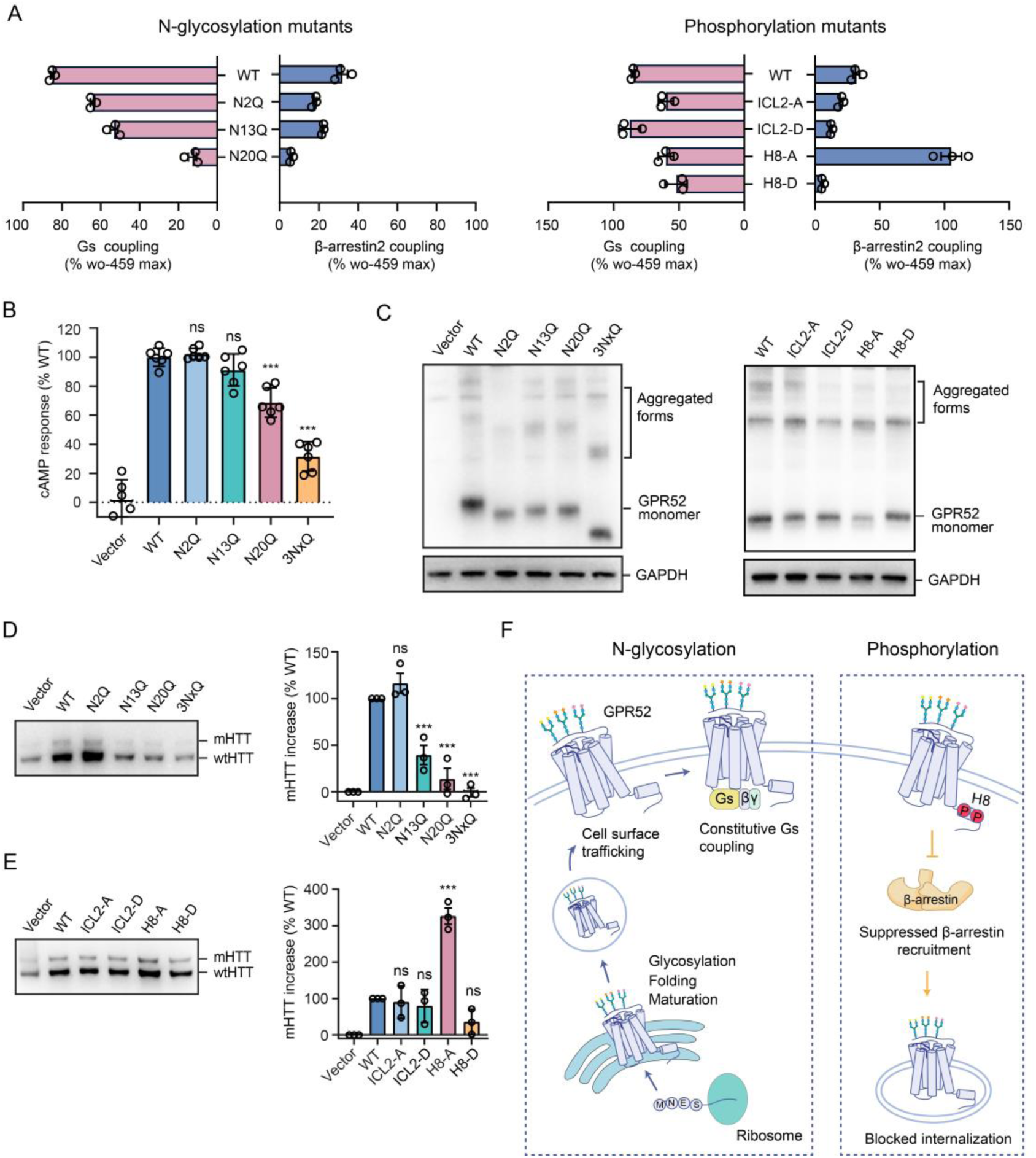
N-glycosylation and phosphorylation orchestrate the intrinsic signaling bias of GPR52 and modulate HTT accumulation in striatal cells. (A) Constitutive Gs *vs* β-arrestin2 coupling profiles of WT and PTM mutants of GPR52. % *E*_max_ is normalized to the agonist-induced full response of WT in the respective pathway. Data are represented as mean ± s.d. from three independent experiments. (B) GPR52-mediated constitutive cAMP accumulation measured in STHdh^Q7/Q111^ cells expressing WT or N-glycosylation mutants. Data are represented as mean ± s.d. from six independent experiments. (C) Western blots of WT, N-glycosylation (left) or phosphorylation (right) mutants of HA-tagged GPR52 expressed in STHdh^Q7/Q111^ cells. GAPDH is the loading control. (D, E) Western blots of HTT levels in STHdh^Q7/Q111^ cells expressing WT, N-glycosylation mutants (D) or phosphorylation mutants (E) of HA-tagged GPR52. Both mHTT and wtHTT levels are detected with Ab1 antibody. Quantification of mHTT levels are shown on the right, and data are represented as mean ± s.e.m from three independent experiments. (F) A proposed model of distinct PTM functions in orchestrating the intrinsic signaling bias of GPR52. Statistics significance was assessed by one-way ANOVA with Dunnett’s multiple comparison test. \**p* < 0.05, \*\**p* < 0.01, \*\*\**p* < 0.001 and ns, no significance.

GPR52 with enriched expression in the brain region striatum is reported to modulate HTT levels and toxicity in multiple HD models^22-24^. To examine the role of PTMs in mediating GPR52 function in striatal neurons, we used the STHdh^Q7/Q111^ cells, a well-established mouse striatal-derived cellular HD knock-in model endogenously expressing both wtHTT and mHTT^24,40^. Heterologous expression of GPR52 is reported to constitutively induce an elevation of both wtHTT and mHTT levels in STHdh^Q7/Q111^ cells with a concomitant induction of cAMP levels^22-24^. In our experiment, endogenous mouse Gpr52 was genetically deleted in these striatal cells before they were transfected with wild-type or mutant human GPR52 (Supplementary Figure 10E). In accordance with our results from HEK239T cells, cAMP signaling was significantly impaired in striatal cells expressing N20Q and 3NxQ mutants, marginally reduced in cells expressing the N13Q mutant, and barely affected in cells expressing the N2Q mutant, though four mutants had similar expression levels (Figures 7B, 7C). Accordingly, lowered HTT levels were observed in striatal cells expressing the N13Q, N20Q, and 3NxQ mutants (Figure 7D), which supports the previous finding that GPR52 modulates HTT accumulation in striatal cells in a cAMP-dependent manner^24^.

Remarkably, we observed a robust increase of both wtHTT and mHTT levels in striatal cells expressing the phosphorylation mutant H8-A whereas expression of other mutants (ICL-A/D and H8-D) had almost no effects on HTT levels relative to wild-type (Figure 7E). As the H8-A mutant exhibits arrestin-biased signaling as opposed to the wild-type, this result indicates that GPR52-mediated HTT accumulation in striatal cells may also depend on arrestin coupling. In summary, while N-glycosylation at N20 could up-regulate HTT levels mainly through Gs-mediated cAMP signaling, phosphorylation in helix 8 could down-regulate HTT levels by suppressing arrestin coupling. We speculate these counteracting effects from two types of PTMs on GPR52 would allow striatal cells to strictly control HTT levels at different physiological conditions.

## Discussion

The ‘signaling bias’ paradigm generally refers to the ligand-dependent activation of certain pathways over others by a single receptor. Intriguingly, our study uncovers a ligand-independent self-driven preference of the Gs pathway over the arrestin pathway by orphan GPR52, which is named the intrinsic signaling bias. Several orphan GPCRs are known to display high basal G protein-coupled activities whereas their arrestin pathway activities have been barely characterized^11-13^. For GPR52, our study indicates its high constitutive Gs coupling does not translate to equally high basal arrestin coupling. Whether such an unbalanced coupling profile exists for other self-activated orphan GPCRs is an interesting question for future investigation.

Previous structural studies of GPR52 and another orphan GPR21 revealed an agonist-like motif in ECL2 that occupies the orthosteric pocket and confers the basal Gs activity for both receptors^11,12^. In line with the structural insight, we found that N-glycosylation at residue N20 is able to promote receptor self-activation and Gs coupling possibly through favoring formation of the ECL2 native fold and Gs binding site in unliganded GPR52 while not altering ligand binding to the allosteric side pocket. In addition, three identified N-glycosylation sites at the N terminus all contribute to the transport of the receptor to the cell surface. Thus, beyond the conventional role of N-glycosylation in GPCR biosynthesis and trafficking, we uncovered a unique role of N-glycosylation in mediating GPCR conformation and constitutive signaling. Further characterization of the interaction between the site-specific N-glycan and ECL2 would shed new light on the regulating mechanism of GPCR N-glycosylation. Profiling the receptor phosphorylation patterns and examining regulatory roles of motif-specific phosphorylation led to novel findings on how this PTM influences the intrinsic signaling bias of GPR52. While phosphorylation in ICL2 plays a conventional role in facilitating arrestin recruitment to GPR52, phosphorylation in helix 8 exerts an opposite effect of suppressing arrestin recruitment to both unliganded and agonist-treated GPR52. GPCR phosphorylation is found to predominantly occur on the C-tail and intracellular loops and plays a major role in promoting arrestin binding and activation^38,41,42^. Previous studies have reported the inhibitory effects of one or two C-tail phosphosites on arrestin binding to different GPCRs by biophysical analyses or MD simulations^42,43^. Aligning with these reports using synthetic C-tail phosphopeptides, our study has provided cell-based pharmacological data with the full-length GPR52 to demonstrate the autoinhibitory role of phosphorylation on helix 8. As indicated by MS quantification, helix 8 has a robust basal phosphorylation abundance especially at S332 such that the inhibitory effect dominates the signaling outcome, resulting in substantially compromised arrestin coupling and blockage of internalization for unliganded GPR52. Importantly, we postulate that the suppression of constitutive arrestin coupling mainly by basal phosphorylation on helix 8 enables GPR52 to maintain its cell surface localization and elicit constant Gs signaling without undergoing spontaneous desensitization and internalization. Consistent with this model, agonist-treated GPR52 reduces its phosphorylation level in helix 8, thus partially relieving the autoinhibition and favoring arrestin recruitment, although the overall receptor internalization is still unappreciable.

The discovery of the inhibitory role of helix 8 phosphorylation in arrestin binding raises a key question in the underlying structural mechanism, considering that helix 8 is not engaged in the major arrestin-binding pocket as revealed by most GPCR-arrestin complex structures^41,44-46^. However, a recent study points out the crucial role of helix 8 in accommodating β-arrestin1 which adopts a special tail-binding conformation^47^. Thus, it remains to be clarified whether phosphorylation in helix 8 of GPR52 suppresses arrestin recruitment by occluding the binding site directly or exploiting an indirect mechanism. It is also worthy of future exploration as to which GRKs or other kinases are responsible for installing the unique phosphorylation pattern on GPR52.

Taken together, we propose a model to summarize distinct functions of PTMs in orchestrating the intrinsic signaling bias of GPR52 as elaborated above (Figure 7F). Moreover, N-glycosylation at the N-terminus and phosphorylation in helix 8 are found to differentially modulate GPR52-dependent HTT accumulation in striatal cells. While PTMs remain unexplored or ignored for many GPCRs, our study demonstrates that a strikingly rich set of new insights can be gained on the molecular mechanism of GPCR activation, signaling, and physiological functions through the lens of PTMs.

## Methods

### Cell culture

HEK293T and STHdh^Q7/Q111^ cells were cultured in Dulbecco’s modified Eagle’s medium, supplemented with 10% fetal bovine serum (FBS, Gibco), 100 IU Penicillin and 100 μg/mL Streptomycin (Gibco). HTLA cells (derived from HEK293 cells with stably expressing a tTA-dependent luciferase reporter and a β-arrestin2-TEV fusion gene) were maintained in DMEM supplemented with 10% FBS (Gibco), 2 μg/mL puromycin (Life) and 100 μg/mL hygromycin B (Mirusbio). HEK293T and HTLA cells were cultured in a humidified atmosphere at 37 ℃ in 5% CO_2_, while STHdh^Q7/Q111^ cells were incubated at 33 ℃ in 5% CO_2_.

### Molecular cloning and mutagenesis

The human full-length GPR52 (residues 1-361) was subcloned into the XbaI and XhoI sites of pcDNA 3.1 vector with a haemagglutinin signal peptide, followed by an HA tag or a FLAG tag at the N terminus using the NovoRec plus One step PCR Cloning kit (Novoprotein). N-glycosylation and phosphorylation mutants were generated using Phanta Max Super-Fidelity DNA Polymerase (Vazyme Biotech Co., Ltd). The constructs for Tango and BRET assays were created as previously described ^26,48^. All the plasmids were verified by DNA sequencing (BioSune).

### Protein expression and purification

The gene of human GPR52 (residues 1-340) with two point mutations (A130^3.41^W and C314^7.50^P) and a 10× His-tag at the C terminus was cloned into a modified pTT5 vector (Invitrogen) ^11^. Haemagglutinin signal peptide, FLAG tag and thermo-stabilized *E. coli* apocytochrome b_562_RIL (BRIL) were inserted into the N terminus of GPR52 to enhance receptor expression and stability. The plasmids encoding wild-type or N-glycosylation mutants of GPR52 were transfected into HEK293F cells (Gibco) cultured at 37 ℃ in 5% CO_2_. Cells were harvested by centrifugation at 48 h post-transfection. The cell membrane isolation and protein purification were performed as previously described^11^. In brief, cell pellets were lysed and washed by repeated washing and ultracentrifugation in the hypotonic buffer of 10 mM HEPES (pH 7.5), 10 mM MgCl_2_, 20 mM KCl, and the high osmotic buffer of 10 mM HEPES (pH 7.5), 1.0 M NaCl, 10 mM MgCl_2_, 20 mM KCl, both containing an EDTA-free protease inhibitor cocktail and a phosphatase inhibitor cocktail (Roche). The membrane pellets were solubilized in the buffer of 50 mM HEPES (pH 7.5), 500 mM NaCl, 1% (w/v) DDM and 0.2% (w/v) CHS for 3 h at 4 ℃. After ultracentrifugation at 36,000 rpm for 30 min at 4 ℃, the supernatant was incubated with TALON IMAC resin (Clontech) at 4 ℃ overnight. The next day, the resin was washed with 10 column volumes of wash buffer (25 mM HEPES, 500 mM NaCl, 0.01% (w/v) DDM, 0.002% (w/v) CHS, 20 mM imidazole, 5% (v/v) glycerol), and proteins were eluted in 5 column volumes of wash buffer with the imidazole concentration at 220 mM. The purity and monodispersity of the GPR52 protein were evaluated by analytical size-exclusion chromatography (aSEC). The protein concentration was determined with the BCA Protein Assay Kit (Tiangen Biotech Co., Ltd).

### PNGase F treatment

HEK293T cells expressing FLAG-tagged GPR52 were harvested and incubated with PNGase F (prepared in-house, 3 μg enzyme for 1 million cells) in 1× Hank’s balanced salt solution (HBSS) (Gibco) buffer at 37 ℃ for 2 h. Then cells were washed with HBSS buffer (Gibco) for three times and processed for cAMP accumulation measurement or western blot analysis. For deglycosylation of purified protein, protein was incubated with PNGase F at a ratio of 1:10 (enzyme to protein) at 4 ℃ overnight prior to in-solution protein digestion.

### Western blot

HEK293T or endogenous mouse Gpr52 knockout STHdh^Q7/Q111^ cells were transfected with wild-type or mutants of GPR52 using the Lipo8000 (Beyotime) transfection reagent. After 24 h (for HEK293T cells) or 48 h (for STHdh^Q7/Q111^ cells) of transfection, cells were lysed in phosphate buffered saline (PBS, Gibco) with 1% Triton X-100 (Roche) containing a protease inhibitor cocktail (Pierce, Thermo) at 4 ℃ for 1 h. Following centrifugation at 16,000*g* for 30 minutes at 4 ℃, the supernatant was transferred to a new microtube. The protein concentration was determined by BCA Protein Assay Kit (Tiangen Biotech Co., Ltd).

For the measurement of HTT accumulation in STHdh^Q7/Q111^ cells, equal amounts of protein samples were loaded and separated on Tris-Acetate, 3-8% gels (BeyoGel, Beyotime). For GPR52 expression measurement, equal amounts of protein samples were loaded and separated on 10% SDS gels (GenScript). Subsequently, the gels were transferred to PVDF membranes (Millipore), blocked by 5% skim milk (Sangon Biotech) for 1 h at room temperature (RT), and then incubated with corresponding antibodies overnight at 4 ℃. The antibodies used in this study included anti-FLAG-M2-HRP (Sigma, A8592, 1:5000), anti-HA (Cell Signaling, C29F4, 1:5000), anti-HTT (Millipore, MAB2166, 1:1000). The HRP-conjugated secondary antibodies used were anti-mouse (CWBIO, CW0102S, 1:5000), anti-rabbit (CWBIO, CW0103S, 1:5000). After washing with TBST, the membranes were detected using an enhanced chemiluminescence kit (Epizyme Biotech) on the imager system (Bio-Rad).

### In-gel digestion and phosphopeptide enrichment for GPR52 overexpressed in cells

The plasmid of full-length GPR52 was transfected into HEK293F cells. After 48 h transfection, cells were harvested and stimulated with or without 10 μM wo-459 for 30 min at 37 ℃ in HBSS buffer. Cell membranes were isolated according to the procedures described in the protein purification section. The membrane pellet was resuspended in the lysis buffer of 4% SDS, 4 M urea and 100 mM Tris-HCl. The resulting membrane protein extract was loaded on a 10% SDS gel (GenScript) for protein separation. Western blot analysis using anti-FLAG was performed in parallel so as to pinpoint the position of GPR52 in the gel. The gel band corresponding to GPR52 was excised and collected for in-gel digestion as previously described with modifications^49^. In brief, the protein band was reduced with 5 mM Tris(2-carboxyethyl)phosphine-hydrochloride (TCEP), and alkylated with 20 mM iodoacetamide (IAA) at RT. Then samples were digested first with chymotrypsin (Promega) at a ratio of 1:80 (enzyme to protein) at RT for 3 h, later with trypsin (Promega) at a ratio of 1:50 (enzyme to protein) at 37 ℃ overnight. Next day, the peptides were extracted with acetonitrile and 0.1% trifluoroacetate (TFA) for three times, and concentrated in a SpeedVac machine (Labconco).

For phosphopeptide enrichment, total peptides from in-gel digestion were re-dissolved with a loading buffer containing 20% lactic acid, 70% ACN and 5% TFA, and then incubated with titanium dioxide beads GL Sciences) at a protein:bead ratio of 5:1 and at RT for 30 min. The beads were pelleted by centrifugation, and then washed with 30% ACN and 80% ACN, both containing 0.5% TFA for 5 times. The phosphopeptides were eluted with 5% NH3·H_2_O (Sigma) twice and lyophilized in a SpeedVac machine (Labconco).

### In-solution digestion of purified GPR52 protein

Purified GPR52 protein was re-solubilized and denatured using 8 M urea, 50 mM Tris-HCl (pH 8.0). Then protein samples were reduced with 5 mM TCEP for 30 min at RT followed by alkylation with 20 mM IAA at RT for another 30 min. Subsequently, samples were digested in solution first with chymotrypsin (Promega) at a ratio of 1:80 (enzyme to protein) at RT for 3 h, later with trypsin (Promega) at a ratio of 1:50 (enzyme to protein) at 37 ℃ overnight. Protease digestion was terminated by adding 1% TFA, and then the peptides were desalted with a C18 microspin column (Omicsolution).

### Mass spectrometry data acquisition

The peptide samples were redissolved with 0.1% FA and analyzed using a nanoElute LC system coupled to a timsTOF Pro 2 mass spectrometer (Bruker). For N-glycosylation data acquisition, peptide samples were separated on an analytical column (75 μm i.d., 300 mm) in-house packed with C18-AQ 1.9-μm C18 resin (Dr. Maisch GmbH) with a gradient of 3–22 % solvent B (0.1% FA in ACN) in 40 min, 22–37% B in 10 min, 37–80% B in 5 min, and 80% B for 5 min at a flow rate of 300 nL/min. DDA (data-dependent acquisition) MS data were acquired in the DDA PASEF mode with the following settings: capillary voltage, 1,500 V; mass range for MS scans, 100 to 1,700 m/z; ion mobility scanning from 0.6 to 1.6 Vs/cm^2^; ramp time, 100 ms; total cycle time, 1.1 s; target intensity (20,000); intensity threshold (2,500); charge range (0-5); isolation width (2 *m/z* for *m/z* < 700 and 3 *m/z* for *m/z* > 700); collisional energy (20-59).

For GRP52 phosphorylation analysis, both unenriched and phospho-enriched samples were separated with a gradient of 3–22 % solvent B (0.1% FA in ACN) in 65 min, 22–37% B in 15 min, 37–80% B in 5 min, and 80% B for 5 min at a flow rate of 300 nL/min. DDA MS data were acquired in the DDA PASEF mode with the following settings: capillary voltage, 1,500 V; mass range for MS scans, 100 to 1,700 m/z; ion mobility scanning from 0.6 to 1.6 Vs/cm^2^; ramp time, 166 ms; total cycle time, 1.9 s; target intensity (20,000); intensity threshold (1,000); charge range (0-5); isolation width (2 *m/z* for *m/z* < 700 and 3 *m/z* for *m/z* > 700); collisional energy (20-59).

### Mass spectrometry data analysis

MS data acquired on timsTOF Pro were processed by FragPipe (version 20.0) integrating MSFragger (version 3.8)^50^. The bulit-in LFQ workflow was employed for glycosylation and phosphorylation site identification. MS data was searched with the following parameters: precursor and fragment mass tolerance 20 ppm; protein digestion with trypsin/chymotrypsin and allowed missed cleavage of 4; carbamidomethylation (C) as fixed modification; oxidation (M) and acetylation (Protein N-term) as variable modifications; phosphorylation (S, T, Y) as variable modification for phosphosite identification; and deamination (N) as variable modification for N-glycosite identification. For all searches, 5 modifications at maximum were allowed. In the validation step, PSM validation and protein inference were processed by Percolator and ProteinProphet integrated in Philosopher (version 5.0.0). In the command line options of PTMProphet, MINPROB indicating the PTM site probability threshold was set to 0.75 and the match-between-run function was enabled. Other settings remained as default. All peptide precursors bearing the same PTM site were clustered and their reported MS intensities were summed for site-specific quantification. For PTM stoichiometry estimation, modified peptide precursors bearing a given PTM site and unmodified counterparts were collected. Then site-specific glycosylation or phosphorylation stoichiometry was represented by the ratio of the summed intensity of modified peptide precursors to that of modified and unmodified precursors.

### LiP-MS experiment and data analysis

About 10 μg purified GPR52 protein for wild-type and N-glycosylation mutants were prepared in a buffer of 25 mM HEPES (pH 7.5), 500 mM NaCl, 0.01% DDM/0.002% CHS, 10% glycerol, 220 mM imidazole, and incubated with Proteinase K (PK, Sigma) at a ratio of 1:50 (enzyme to protein) at 25 ℃ for 5 min for limited proteolysis. Protein samples were then heated at 98 ℃ for 10 min to terminate proteolysis. After lyophilization in a SpeedVac concentrator, samples were re-solubilized and denatured using a solution of 5% SDC and 4 M urea. Denatured samples were then reduced by 5 mM TCEP for 30 min at RT and alkylated by 20 mM IAA for 30 min at RT in the dark. Samples were diluted with 100 mM ammonium bicarbonate to a final concentration of 1.25% SDC and 1 M urea. For complete proteolysis, protein samples were then incubated with chymotrypsin at a ratio of 1:80 (enzyme to protein) for 3-4 h at RT and then trypsin (Promega) at a ratio of 1:80 (enzyme to protein) overnight at 37 ℃. After quenching the digestion with 1% TFA, peptide samples were desalted by a C18 microspin column (Omicsolution) and lyophilized in a SpeedVac concentrator. For GPR52 wild-type or mutant proteins, their LiP-MS samples were all prepared in four independent replicates. Additionally, we prepared a wild-type complete digestion sample in quadruplicate using chymotrypsin and trypsin without PK treatment, which was used to generate a reference tryptic peptide list for LiP-MS data analysis.

Peptide samples were re-dissolved with 0.1% FA and analyzed using an EASY-nLC 1200 system (Thermo Fisher Scientific) coupled to a Q-Exactive HF mass spectrometer (Thermo Fisher Scientific). The peptides were separated on an analytical column (75 μm i.d., 200 mm) in-house packed with C18-AQ 1.9-μm C18 resin (Dr. Maisch GmbH) using a gradient of 3 to 5% solvent B (0.1% FA in acetonitrile) in 13 min, 5-21% B in 46 min, 21-45% B in 21 min, 45-100% B in 4 min and 100% B for 6 min at a flow rate of 300 nL/min. For LiP-MS samples from the wild-type and each mutant protein, DDA and DIA data were acquired for each replicate. For data acquisition in the DDA mode, MS2 spectra were generated for up to 15 peptide precursors with an isolation window of 1.4 *m/z*, and fragments were detected at a resolution of 15,000. The MS2 AGC target value was set to 1 × 10^5^ with a maximum ion injection time of 45 ms in MS2. For data acquisition in the DIA mode, the precursor ions were fragmented in 22 variable windows covering a range of 350 to 1200 m/z. The resolution of Orbitrap analyzer was set to 60,000 for MS1 and 30,000 for MS2. The AGC was set to 3 × 10^6^ in MS1 and 1 × 10^6^ in MS2 with a maximum injection time was set to 20 ms in MS1 and 50 ms in MS2. The NCE was set to 28%.

LiP-MS data was then processed using Spectronaut (version 17.3, Biognosys). We first generated a project-specific spectral library by importing all DDA raw files into Spectronaut with default settings. Trypsin and chymotrypsin were set as specific enzymes, cabamidomethyl (C) was set to fixed modification, and oxidation (M) and acetyl (protein N-terminus) were set to variable modifications. Maximal four missed cleavages were allowed and FDRs on PSM/peptide/protein levels were all set to 1%. Then DIA data from all LiP-MS samples were combined and processed using the previously built spectral library in Spectronaut with default settings: Q value of precursor and protein cutoff set to 0.01; decoy generation method “Mutated”; quantification based on MS2 area; normalization strategy set to “false”. The peptide quantification reports were exported, and only those present in the tryptic peptide reference list were retained for further analysis. In the pairwise comparison of each mutant *vs* wild-type GPR52, quantified tryptic peptides with an average fold-change >2 or <0.5 (*p* <0.05, n=4) or peptides exclusively detected in wild-type or mutant samples in at least three replicates are defined as differential peptides according to the general criteria for differential analysis in LiP-MS^28,30^. Statistical significance for differential peptides was assessed using the limma package (version 3.59.10) in R (version 4.4.0). Structural regions covered by quantified tryptic peptides in LiP-MS analysis are color-coded in the GPR52 snake plot yielded by GPCRdb^51^ to highlight structurally altered regions in each mutant relative to wild-type.

### cAMP accumulation assay

Intracellular cAMP accumulation was measured using the Homogeneous Time-Resolved Fluorescence (HTRF) kit (Cisbio) according to the manufacture’s introduction. In brief, HEK293T or STHdh^Q7/Q111^ cells were transfected with the indicated receptor constructs using Lipo8000 transfection reagent (Beyotime). After 24 h (for HEK293T cells) or 48 h (for STHdh^Q7/Q111^ cells) transfection, cells were detached from the plate with cell dissociation buffer (Sigma) and collected into a new tube by centrifugation. Sequentially, the cells were suspended in DMEM containing 500 μM IBMX (Sigma) and transferred into a white 384-well plate (Thermo Fisher) at a density of 2000 cells per well. Cells were treated with 5 μL agonist at various concentrations (2×) (for agonist concentration-response measurement) or 5 μL assay buffer (for constitutive activity measurement) for 30 min. Then, cells were incubated with cAMP-D2 and anti-cAMP Crypate (Cisbio) for 1 h at RT. The plate was read on an Envision reader (Perkin Elmer). For determination of agonist concentration-response curves, data were normalized to percent wild-type signals and EC_50_ was analyzed in GraphPad Prism 9.0 (Graphpad) using log (agonist) vs response. For measurement of receptor constitutive activities, data were normalized to the baseline signals of the vector (set to 0%) and to the maximum signals induced by agonists (set to 100%).

### Tango assay for β-arrestin2 recruitment measurement

Tango assay was performed to measure the β-arrestin2 recruitment activity of GPR52 following a previously described protocol^48^. Briefly, HTLA cells were transfected with Tango plasmids encoding wild-type and mutants of GPR52 using the calcium phosphate method. After 24 h transfection, HTLA cells were plated into poly-lysine coated 384-well white clear bottom plates at a density of 10,000-15,000 cells per well with 40 μL DMEM containing 1% dialyzed FBS (Gibco). Six hours later, cells were treated with 20 μL agonist at various concentrations (3×) (for agonist concentration-response measurement) or 20 μL assay buffer (for constitutive activity measurement). The plates were then incubated at 37 °C overnight. The next day, culture media were decanted and 20 μL BrightGlo reagent (Promega) was added into each well. After 20 min incubation in the dark at RT, luminescence was read on an Envision counter (Perkin Elmer). For determination of agonist concentration-response curves, data were normalized to percent wild-type signals and EC_50_ was analyzed in GraphPad Prism 9.0 (Graphpad) using log (agonist) vs response. For measurement of receptor constitutive activities, data were normalized to the baseline signals of the vector (set to 0%) and to the maximum signals induced by agonists (set to 100%).

### BRET assays for Gs dissociation and β-arrestin recruitment measurement

BRET-based Gs dissociation assays were performed using the published TRUPATH system as previously described^26^. Briefly, HEK293T cells were transfected with receptor, Gαs-RLuc8, Gβ1 and Gγ1-GFP2 at a ratio of 1:1:1:1 (100 ng receptor per well for a 6-well plate) using the calcium phosphate method. After 24 h transfection, cells were seeded into a poly-D-lysine-coated white 96-well assay plate (Corning) at a density of 50,000 cells per well and incubated at 37 °C overnight. The next day, culture media was aspirated and replaced with 40 μL of assay buffer (1× HBSS + 20 mM HEPES, pH 7.4) containing 4 μM freshly prepared coelenterazine 400a (Nanolight Technologies) in the dark at RT for 5 min. For the basal Gs dissociation activity measurement, the plate was read in an LB940 Mithras plate reader (Berthold Technologies) with 410 and 515 nm emission filters. For the agonist-induced Gs dissociation activity assessment, cells were stimulated with 20 μL agonist at various concentrations (3×) before reading. BRET2 ratios were calculated as the ratio of the GPF2 to RLuc8 signals.

For BRET-based β-arrestin (β-arrestin1 and β-arrestin2) recruitment assays, we used the receptor-RLuc8 and β-arrestin-GFP2 as BRET biosensors. HEK293T cells were transfected with receptor-RLuc8 and β-arrestin-GFP2 at a ratio of 1:4 (150 ng receptor-RLuc8 per well for a 6-well plate) using the calcium phosphate method. After 24 h transfection, cells were plated into a poly-D-lysine-coated white 96-well assay plate (Corning) at a density of 50,000 cells per well and incubated at 37 °C overnight. The following day, after aspiration of the media, cells were incubated with 40 μL of assay buffer (1× HBSS + 20 mM HEPES, pH 7.4) containing 4 μM freshly prepared coelenterazine 400a (Nanolight Technologies) at RT for 5 min. Then, the plate was read in an LB940 Mithras plate reader (Berthold Technologies) with 410 and 515 nm emission filters. For agonist-induced β-arrestin recruitment activity measurement, cells were stimulated with 40 μL various concentrations (2×) of agonist for 10 min prior to the addition of coelenterazine 400a. BRET2 ratios were calculated as the ratio of the GPF2 to RLuc8 signals.

For determination of agonist concentration-response curves, data were normalized to percent wild-type signals and EC_50_ was analyzed in GraphPad Prism 9.0 (Graphpad) using log (agonist) vs response. For measurement of receptor constitutive activities, data were normalized to the baseline signals of the vector (set to 0%) and to the maximum signals induced by agonists (set to 100%).

### Receptor internalization assay by FACS

The internalization assay was performed with a previously described method^52^. In brief, 3 μg plasmids of HA-tagged wild-type or phosphorylation mutants were transfected into HEK293T cells cultured in 10 cm dishes using the calcium phosphate method. After 24 h transfection, cells were detached with the cell dissociation buffer (Sigma) and washed with HBSS for twice by centrifugation. Cells were suspended in HBSS buffer and chilled on ice for 10 min. Then cells were incubated with 10 μM wo-459 for 0, 5, 10, 15, 30, 60, 120 and 240 min at 37 °C. After agonist stimulation, cells were fixed with 4% paraformaldehyde (PFA) at RT for 20 min. Cells were blocked with HBSS buffer containing 3% BSA for 20 min followed by incubation with monoclonal anti-HA-FITC antibody (1:500) (Sigma, H7411) for 1 h at RT in the dark. Then cells were washed twice with PBS and analyzed on a CytoFLEX flow cytometry (Beckman). Fluorescent signal from a FITC-conjugated anti-HA antibody was recorded in a 488 nm channel and flow cytometry data were analyzed by Flow Jo 10.4 software (BD Biosciences). Untransfected cells were used to gate cells with specific fluorescence signals. Mean fluorescence intensities were used for quantification. Data were normalized to the signals detected from cells without agonist stimulation.

### Receptor internalization assay by immunofluorescence staining

To visualize the internalization of GLP-1R and GPR52, HEK293T cells were plated on coverslips coated with 0.5% Matrigel (Corning) and transfected with FLAG-tagged receptor plasmids for 24 h. Cells were incubated with a primary antibody (mouse anti-FLAG-M2, Sigma, F1804, 1:500) at 4 °C for 1 h in HBSS buffer. Cells were washed with chilled HBSS buffer for three times and then incubated with or without an agonist at 37 °C for different time points followed by fixation with 4% PFA for 20 min at RT. Cells were blocked with 3% BSA in HBSS buffer for 20 min and permeabilized with 0.1% Triton X-100 (Roche) in blocking buffer for another 20 min at RT. Cells were then incubated with secondary antibody (Alexa-488-conjugated goat anti-mouse, Abcam, ab150113, 1:500) for 1 h at RT and mounted on glass slides using ProLong Gold Antifade mounting medium (Invitrogen). Fluorescence confocal microscopy was performed with a laser scanning confocal microscope (Nikon A1R) with a 100× oil immersion objective.

To monitor the internalization of wild-type or phosphorylation mutants of GPR52, HA-tagged receptor plasmids were transfected into HEK293T cells on coverslips. After 24 h transfection, cells were incubated with 10 μM wo-459 in HBSS buffer for different time points at 37 °C followed by fixation with 4% PFA for 20 min at RT. Cells were blocked with 3% BSA in HBSS buffer for 20 min and incubated with a primary antibody (rat anti-HA, Roche, RD11867423001, 1:100) overnight at 4 °C. Then cells were washed three times with blocking buffer before incubation with a secondary antibody (Alexa-568-conjugated donkey anti-rat, Invitrogen, A78946, 1:500) at RT for 1 h. Coverslips were mounted on glass slides using ProLong Gold Antifade mounting medium (Invitrogen). Images were collected on a spinning disk microscope (Nikon) with a 100× oil immersion objective.

Image analysis was carried out using Fiji (v1.54f) and mean intensities of the region of interest (ROI) draw around receptor-positive cells (based on 488 nm or 568 nm fluorescent signals) were measured for quantification. Data were normalized to the signals detected from cells without agonist stimulation.

### Cell surface and total expression measurement

The surface and total expression levels of wild-type and mutants of GPR52 were determined by flow cytometry with a procedure very similar to the receptor internalization assay. Briefly, HTLA or HEK293T cells were transfected with FLAG-tagged or HA-tagged plasmids encoding wild-type or individual mutants of GPR52. After 24 h transfection, cells were collected and fixed with 4% PFA for 20 min at RT. Then cells were blocked with 3% BSA for 20 min at RT before incubation with a monoclonal anti-FLAG-M2-FITC (Sigma, F4049, 1:500) or anti-HA-FITC antibody (Sigma, H7411, 1:500) in PBS at RT for 1 h in the dark. For total expression measurement, blocked cells were incubated with a FITC-conjugated anti-FLAG or anti-HA antibody in PBS buffer containing 0.1% Triton X-100 (Roche) for 1 h. Cells were washed with PBS for twice, and then analyzed by flow cytometry. Data were collected on a CytoFLEX Platform (Beckman) and analyzed with Flow Jo 10.4 software (BD Biosciences). Data were normalized to the percentage of wild-type signals.

### Generation of a knockout cell line

Gpr52 knockout STHdh^Q7/111^ cell line was obtained by single cell flow sorting with BD FACSAria™ III Cell Sorter. In brief, 3 µg Cas9 vector and 1 µg individual gRNA vector were simultaneously transfected with Lipofectamine™ 3000 (ThermoFisher) into STHdh^Q7/111^ cells cultured in a 6-well plate (Corning) with 30% cell density. Then flow sorting was performed with GFP located in gRNA vector two days after transfection, and cultured with caspase inhibitor Z-VAD-FMK (25 µM) (MCE) to increase the cell availability. Single cell clones were collected and cultured in 96-well plate (Corning) with flow sorting, and enriched to identify the genotype with Sanger sequencing. gRNAs used in this study were 5’-CGTTGGAGTTACCTGCTTGG-3’ and 5’-GCCTCTATCCTACAATCAAC-3’. Primers used for genotyping were 5’-CCTGCCCTCTTGGATTTGGT-3’ and 5’-TGACTAGGGAATCTGGCCCT-3’.

## Acknowledgments

We thank P. Si at the cell expression core and other staff members at the purification or assay core facilities of iHuman Institute for their technical support. This work was funded by the National Key R&D Program of China (2022YFA1302902), National Natural Science Foundation of China (32171439 to W.S., 32301250 to B.Z., 92049301 to B.L., 81925012 to B.L.), Shanghai Frontiers Science Center for Biomacromolecules and Precision Medicine at ShanghaiTech University, Innovation Program of Shanghai Municipal Education Commission (2021-01-07-00-07.E00074 to B.L.), and the New Cornerstone Science Foundation (NCI202242 to B.L.). We also thank the Shanghai Municipal Government and ShanghaiTech University for financial support.

## Author contributions

W.S. conceived and supervised the project. B.Z. performed cellular activity assays and data analysis. M.M. and W.G. performed protein purification, MS analysis of PTMs and HTT accumulation assay. S.L. performed LiP-MS analysis. G.Y. and H.W. generated the GPR52 KO cell line. B.L. co-supervised the project, made fruitful discussion, and edited the manuscript. W.S. and B.Z. wrote the manuscript with input from all authors.

## Competing Interests

The authors declare no competing financial interests.

## Notes

### Competing Interest Statement

The authors have declared no competing interest.

## References

1. Kolb, P. et al. Community guidelines for GPCR ligand bias: IUPHAR review 32. British Journal of Pharmacology 179, 3651–3674 (2022).

2. Wacker, D., Stevens, R.C. & Roth, B.L. How Ligands Illuminate GPCR Molecular Pharmacology. Cell 170, 414–427 (2017).

3. Hilger, D., Masureel, M. & Kobilka, B.K. Structure and dynamics of GPCR signaling complexes. Nat Struct Mol Biol 25, 4–12 (2018).

4. Wootten, D., Christopoulos, A., Marti-Solano, M., Babu, M.M. & Sexton, P.M. Mechanisms of signalling and biased agonism in G protein-coupled receptors. Nat Rev Mol Cell Biol 19, 638–653 (2018).

5. Che, T. & Roth, B.L. Molecular basis of opioid receptor signaling. Cell 186, 5203–5219 (2023).

6. Manglik, A. et al. Structure-based discovery of opioid analgesics with reduced side effects. Nature 537, 185–190 (2016).

7. Roth, B.L. Molecular pharmacology of metabotropic receptors targeted by neuropsychiatric drugs. Nat Struct Mol Biol 26, 535–544 (2019).

8. Cao, D. et al. Structure-based discovery of nonhallucinogenic psychedelic analogs. Science 375, 403–411 (2022).

9. Schmid, C.L. et al. Bias Factor and Therapeutic Window Correlate to Predict Safer Opioid Analgesics. Cell 171, 1165–1175 e13 (2017).

10. Martin, A.L., Steurer, M.A. & Aronstam, R.S. Constitutive Activity among Orphan Class-A G Protein Coupled Receptors. PLoS One 10, e0138463 (2015).

11. Lin, X. et al. Structural basis of ligand recognition and self-activation of orphan GPR52. Nature 579, 152–157 (2020).

12. Lin, X. et al. Cryo-EM structures of orphan GPR21 signaling complexes. Nat Commun 14, 216 (2023).

13. Chen, G. et al. Structural and functional characterization of the endogenous agonist for orphan receptor GPR3. Cell Res 34, 262–265 (2024).

14. Xiong, Y. et al. Identification of oleic acid as an endogenous ligand of GPR3. Cell Res 34, 232–244 (2024).

15. Qu, X. et al. Structural basis of tethered agonism of the adhesion GPCRs ADGRD1 and ADGRF1. Nature 604, 779–785 (2022).

16. Patwardhan, A., Cheng, N. & Trejo, J. Post-Translational Modifications of G Protein-Coupled Receptors Control Cellular Signaling Dynamics in Space and Time. Pharmacol Rev 73, 120–151 (2021).

17. Zhang, B., Li, S. & Shui, W. Post-Translational Modifications of G Protein-Coupled Receptors Revealed by Proteomics and Structural Biology. Front Chem 10, 843502 (2022).

18. Goth, C.K., Petaja-Repo, U.E. & Rosenkilde, M.M. G Protein-Coupled Receptors in the Sweet Spot: Glycosylation and other Post-translational Modifications. ACS Pharmacol Transl Sci 3, 237–245 (2020).

19. Min, C. et al. N-linked Glycosylation on the N-terminus of the dopamine D2 and D3 receptors determines receptor association with specific microdomains in the plasma membrane. Biochim Biophys Acta 1853, 41–51 (2015).

20. Virion, Z. et al. Sialic acid mediated mechanical activation of beta(2) adrenergic receptors by bacterial pili. Nat Commun 10, 4752 (2019).

21. Ali, S., Wang, P., Murphy, R.E., Allen, J.A. & Zhou, J. Orphan GPR52 as an emerging neurotherapeutic target. Drug Discov Today 29, 103922 (2024).

22. Wang, C. et al. GPR52 Antagonist Reduces Huntingtin Levels and Ameliorates Huntington’s Disease-Related Phenotypes. J Med Chem 64, 941–957 (2021).

23. Song, H. et al. Targeting Gpr52 lowers mutant HTT levels and rescues Huntington’s disease-associated phenotypes. Brain 141, 1782–1798 (2018).

24. Yao, Y. et al. A striatal-enriched intronic GPCR modulates huntingtin levels and toxicity. Elife 4(2015).

25. Cary, B.P. et al. Structural and functional diversity among agonist-bound states of the GLP-1 receptor. Nat Chem Biol 18, 256–263 (2022).

26. Olsen, R.H.J. et al. TRUPATH, an open-source biosensor platform for interrogating the GPCR transducerome. Nat Chem Biol 16, 841–849 (2020).

27. de Souza, N. & Picotti, P. Mass spectrometry analysis of the structural proteome. Curr Opin Struct Biol 60, 57–65 (2020).

28. Malinovska, L. et al. Proteome-wide structural changes measured with limited proteolysis-mass spectrometry: an advanced protocol for high-throughput applications. Nat Protoc 18, 659–682 (2023).

29. Cappelletti, V. et al. Dynamic 3D proteomes reveal protein functional alterations at high resolution in situ. Cell 184, 545–559 e22 (2021).

30. Mackmull, M.T. et al. Global, in situ analysis of the structural proteome in individuals with Parkinson’s disease to identify a new class of biomarker. Nat Struct Mol Biol 29, 978–989 (2022).

31. Zhang, B. et al. A Novel G Protein-Biased and Subtype-Selective Agonist for a G Protein-Coupled Receptor Discovered from Screening Herbal Extracts. ACS Cent Sci 6, 213–225 (2020).

32. Xin, Y. et al. Affinity selection of double-click triazole libraries for rapid discovery of allosteric modulators for GLP-1 receptor. Proc Natl Acad Sci U S A 120, e2220767120 (2023).

33. Lu, Y. et al. Accelerating the Throughput of Affinity Mass Spectrometry-Based Ligand Screening toward a G Protein-Coupled Receptor. Anal Chem 91, 8162–8169 (2019).

34. Ma, M. et al. Targeted Proteomics Combined with Affinity Mass Spectrometry Analysis Reveals Antagonist E7 Acts As an Intracellular Covalent Ligand of Orphan Receptor GPR52. ACS Chem Biol 15, 3275–3284 (2020).

35. Maharana, J., Banerjee, R., Yadav, M.K., Sarma, P. & Shukla, A.K. Emerging structural insights into GPCR-beta-arrestin interaction and functional outcomes. Curr Opin Struct Biol 75, 102406 (2022).

36. Dwivedi-Agnihotri, H. et al. Distinct phosphorylation sites in a prototypical GPCR differently orchestrate beta-arrestin interaction, trafficking, and signaling. Sci Adv 6(2020).

37. He, Q.T. et al. Structural studies of phosphorylation-dependent interactions between the V2R receptor and arrestin-2. Nat Commun 12, 2396 (2021).

38. Yang, Z. et al. Phosphorylation of G Protein-Coupled Receptors: From the Barcode Hypothesis to the Flute Model. Mol Pharmacol 92, 201–210 (2017).

39. Hatzipantelis, C.J., Lu, Y., Spark, D.L., Langmead, C.J. & Stewart, G.D. beta- Arrestin-2-Dependent Mechanism of GPR52 Signaling in Frontal Cortical Neurons. ACS Chem Neurosci 11, 2077–2084 (2020).

40. Trettel, F. et al. Dominant phenotypes produced by the HD mutation in STHdh(Q111) striatal cells. Hum Mol Genet 9, 2799–809 (2000).

41. Zhou, X.E. et al. Identification of Phosphorylation Codes for Arrestin Recruitment by G Protein-Coupled Receptors. Cell 170, 457–469 e13 (2017).

42. Latorraca, N.R. et al. How GPCR Phosphorylation Patterns Orchestrate Arrestin-Mediated Signaling. Cell 183, 1813–1825 e18 (2020).

43. Mayer, D. et al. Distinct G protein-coupled receptor phosphorylation motifs modulate arrestin affinity and activation and global conformation. Nat Commun 10, 1261 (2019).

44. Huang, W. et al. Structure of the neurotensin receptor 1 in complex with beta-arrestin 1. Nature 579, 303–308 (2020).

45. Staus, D.P. et al. Structure of the M2 muscarinic receptor-beta-arrestin complex in a lipid nanodisc. Nature 579, 297–302 (2020).

46. Cao, C. et al. Signaling snapshots of a serotonin receptor activated by the prototypical psychedelic LSD. Neuron 110, 3154–3167 e7 (2022).

47. Chen, K. et al. Tail engagement of arrestin at the glucagon receptor. Nature 620, 904–910 (2023).

48. Kroeze, W.K. et al. PRESTO-Tango as an open-source resource for interrogation of the druggable human GPCRome. Nat Struct Mol Biol 22, 362–9 (2015).

49. Shevchenko, A., Tomas, H., Havlis, J., Olsen, J.V. & Mann, M. In-gel digestion for mass spectrometric characterization of proteins and proteomes. Nat Protoc 1, 2856–60 (2006).

50. Demichev, V. et al. dia-PASEF data analysis using FragPipe and DIA-NN for deep proteomics of low sample amounts. Nat Commun 13, 3944 (2022).

51. Pandy-Szekeres, G., et al. GPCRdb in 2023: state-specific structure models using AlphaFold2 and new ligand resources. Nucleic Acids Res 51, D395–D402 (2023).

52. Hendrik Schmidt, J., et al. Constitutive internalization across therapeutically targeted GPCRs correlates with constitutive activity. Basic Clin Pharmacol Toxicol 126 Suppl 6, 116–121 (2020).

53. Deutsch, E.W. et al. The ProteomeXchange consortium at 10 years: 2023 update. Nucleic Acids Res 51, D1539–D1548 (2023).

54. Chen, T., et al. iProX in 2021: connecting proteomics data sharing with big data. Nucleic Acids Res 50, D1522–D1527 (2022).

